# Polarity-driven three-dimensional spontaneous rotation of a cell doublet

**DOI:** 10.1101/2022.12.21.521355

**Authors:** Linjie Lu, Tristan Guyomar, Quentin Vagne, Rémi Berthoz, Alejandro Torres-Sánchez, Michèle Lieb, Cecilie Martin-Lemaitre, Kobus van Unen, Alf Honigmann, Olivier Pertz, Guillaume Salbreux, Daniel Riveline

## Abstract

Cell mechanical interactions play a fundamental role in the self-organisation of organisms. How these interactions drive coordinated cell movement in three-dimensions remains unclear. Here we report that cell doublets embedded in a 3D extracellular matrix undergo spontaneous rotations and we investigate the rotation mechanism using live cell imaging, quantitative measurements, mechanical perturbations, and theory. We find that rotation is driven by a polarized distribution of myosin within cell cortices. The mismatched orientation of this polarized distribution breaks the doublet mirror symmetry. In addition, cells adhere at their interface through adherens junctions and with the extracellular matrix through focal contacts near myosin clusters. Using a physical theory describing the doublet as two interacting active surfaces, we find that rotation is driven by myosin-generated gradients of active tension, whose profiles are dictated by interacting cell polarity axes. We show that interface three-dimensional shapes can be understood from the Curie principle: shapes symmetries are related to broken symmetries of myosin distribution in cortices. To test for the rotation mechanism, we suppress myosin clusters using laser ablation and we generate new myosin clusters by optogenetics. Our work clarifies how polarity-oriented active mechanical forces drive collective cell motion in three dimensions.

## Spontaneous rotations *in vivo* and *in vitro*

Spontaneous cell rotational motions have been reported in a variety of contexts *in vivo*. Tissues undergo rotation during development in *Drosophila* in the egg chamber^1^, in the ommatidia of the retina^2^, in the testis^3^, and in zebrafish embryos, where rotation of cell pairs occurs in the zebrafish’s lateral line^4^. In early *C. elegans* embryo development, chiral counter-rotating flows break chiral symmetry and play a role in setting the organism’s left-right axis^5,6^.

Seminal observations *in vitro* in two-dimensions have shown that endothelial adhering cells migrating on a substrate and confined within a two-dimensional pattern form a stably rotating doublet^7^. The cell-cell interface adopts a curved shape, such that the doublet acquires an overall shape reminiscent of a “yin-yang”. More recently, groups of epithelial cells were reported to undergo rotation within rings^8,9^. In three-dimensions *in vitro*, during alveologenesis of the human mammary gland, it was shown that organoids also undergo rotation^10^. In addition, MDCK cells can assemble into hollow cysts in three-dimensions which undergo spontaneous rotation in an assay within two layers of Matrigel^11^. There the two layers of Matrigel impose a polarization axis to the cyst which allows to probe for chiral broken symmetry, revealed in a bias in the direction of rotation. Altogether, rotational flow appears to be a common feature of the collective motion of interacting cells.

However, it is unclear how these rotational movements arise from the distribution of force-generating elements in the cell. Several models have been proposed to explain the rotation of a cell doublet on a two-dimensional substrate confined in a micropattern, using phase-field, particle-based, or cellular Potts models^12–15^. These models exhibit simultaneous doublet rotation and interface deformation, based on a representation of actin polymerization forces and protrusion-forming forces^12–15^ and coupling to a biochemical system exhibiting spontaneous polarization through feedback between an activator and inhibitor^13^, or directly to an internal polarization vector^14^. Despite these advances, it is still unclear what biophysical mechanisms underlie collective cell rotation in three dimensions, and notably how force-generating elements in the cell self-organize to drive coherent cell motion.

## Dynamics of MDCK doublet rotation

Here, we exploit a novel assay to study the mechanism behind the spontaneous rotation in three dimensions of MDCK cell doublets. We confined cells within a thin layer of Matrigel, close to the coverslip, to optimize imaging resolution (Fig. 1a, Methods). Strikingly, all embedded MDCK cell clusters undergo spontaneous rotation. We focus here on adhering cell doublets which emerge from the division of a single cell. There was no obvious common orientation of the axis of rotation of cell doublets, and in most but not all cases, a lumen at the cell-cell interface rotated with the doublet (Fig. 1b, Supplementary Videos 1,2). Single cells instead did not rotate in Matrigel (Ext. Fig. 1). Cells participating in the rotating doublet however do not have to be sister cells, as two cells with different fluorescent E-cadherin labels could adhere to each other and initiate rotation (Fig. 1c and Supplementary Video 3).

**Figure 1.**
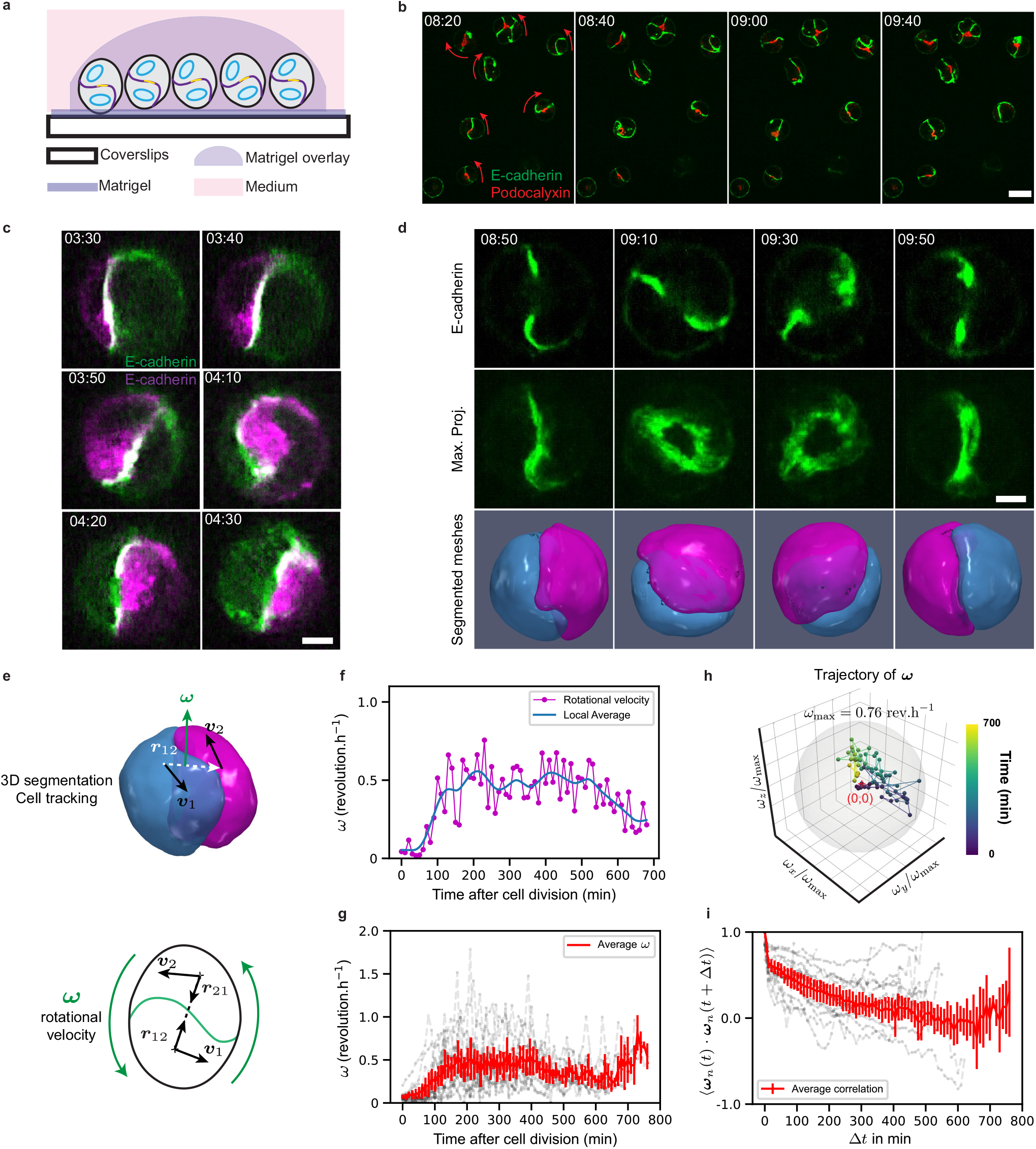
3D rotation of MDCK cell doublet. **a**. Schematics of the experimental assay. **b**. Snapshots of rotating doublets. Red arrows indicate the direction of rotation. Labels: E-cadherin-mNG (Green), Podocalyxin-mScarlet (Red). Time relative to beginning of Movie 1. **c**. Snapshot of a rotating doublet with two cells expressing E-cadherin of different colours. Labels: E-cadherin-GFP (Green) and E-Cadherin-DsRed (Magenta). Time relative to beginning of Movie 3. **d**. Snapshots of a rotating doublet with labelled E-cadherin-mNG (Green) (from top to bottom, cross-section, maximum projection, and three-dimensional segmentation). *n* = 14 doublets, *N* = 3 biological repeats. Time relative to cell division. **e**. Schematics for the calculation of the rotation vector ***ω*** (see Supplementary Information, section 2). **f.** Magnitude of the rotational velocity as a function of time after cell division. **g**. Average and individual trajectories of the magnitude of the rotational velocity of cell doublets after cell division. *n* = 14, *N* = 3. **h**. Trajectory of rotation vector normalized with respect to its maximum amplitude, colormap indicates time from dark blue to yellow. Grey sphere has unit radius. **i**. Autocorrelation of *ω*_n_=***ω***/||***ω***|| as a function of lag time. Scale bars: 5 *μ*m except panel b: 20 *μ*m. Time in panels b,c,d in hh:mm. Error bars: 95% confidence interval of the mean.

To investigate quantitatively doublet rotation (Supplementary Information sections 1, 2), we imaged rotating doublets expressing E-cadherin and computationally segmented cell shapes in the doublet (Fig. 1d, Supplementary Video 4 and Methods). We calculated the center of mass and velocity of each doublet cell, which allows defining a doublet rotation vector ***ω*** (Fig. 1e). The norm of the rotation vector increased after cell division for ~100 minutes, before reaching a roughly constant rotational velocity of ~ 180 degrees/h for a duration of ~10 hours, corresponding to about 5 continuous full turns along the same direction (Fig. 1f, g). This is consistent with previously reported rotation velocities of MDCK cysts and breast epithelial cell spheres^11,16^. Plotting the trajectory of the vector ***ω*** showed that the axis of rotation is not fixed but appears to drift over time (Fig. 1h and Ext. Fig. 2). This is expected to the extent that no external cue sets a preferred axis of rotation. We note that this also implies that the spinning motion of the doublet does not intrinsically break chiral symmetry. Plotting the correlation function of the normalized rotation vector further indicated a characteristic correlation time of a few hours (Fig. 1i).

## Dynamics of doublet elongation during rotation

We then wondered whether doublet cells were rotating relative to each other or were rotating together as a solid object (Fig. 2a). We reasoned that in the latter case, the doublet elongation axis would rotate together with the rotation vector ***ω***. We extracted from computationally segmented doublet shapes (Fig. 1d) the three orthogonal principal axes of elongation and evaluated the corresponding relative elongation magnitudes (Fig. 2b, Supplementary Information section 3). We noted that the elongation of the doublet major axis was maximal at cell division and then relaxed to a nearly constant value, following the same trend as the magnitude of rotation, but in reverse (Fig. 1h, 2b). We then tested whether the rotation axis was correlated with the axis of maximal doublet elongation. This revealed a strong anti-correlation between the major axis of elongation and the direction of rotation (Fig. 2c), indicating that the elongation major axis is within the plane of rotation. The two minor cell elongation axes had instead weak positive correlations to the axis of rotation. The elongation major axis was also aligned with the vector joining the center of mass of the cells, indicating that it rotates together with the double (Fig. 2c). Overall, these results suggest that the doublet rotates as a single physical object. Consistent with this idea, patterns of E-cadherin (Ext. Fig. 3a–b and Supplementary Information section 6, Supplementary Video 4) and actin (Supplementary Video 5) at the cell-cell contact remained similar during ~ 1 hour 30 min of observation.

**Figure 2.**
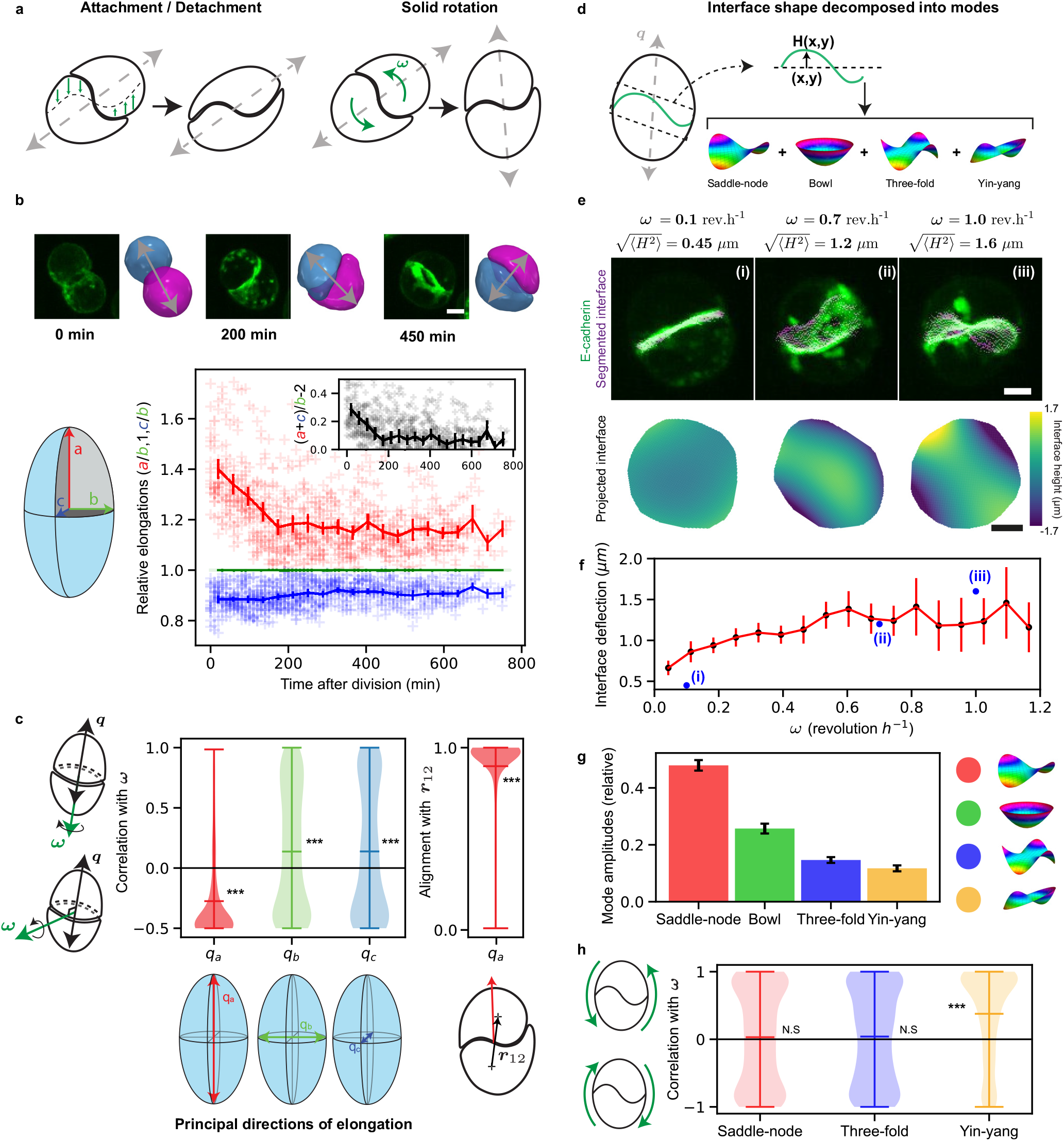
Coordinated rotation of doublet shape and interface. **a**. Schematics of possible scenarios of doublet rotation. Left: The interface is deforming (green arrows), leading to apparent rotation without motion of the doublet outer surface. Right: the doublet is rotating as a solid object and the cell elongation axis rotates with the doublet. **b**. Top: snapshots of rotating doublets with labelled E-cadherin-mNG (Green) and corresponding segmented meshes. Grey double arrows, approximate elongation axis. Bottom left: schematics for definition of three doublet elongation axes. Bottom right: relative doublet elongation magnitudes as a function of time after division. Inset: ratio of elongation magnitudes, positive values indicate a prolate shape. **c**. (Left) Correlation of axis of rotation with axes of elongation, as indicated in the schematics. The direction of maximal elongation, *q_a_* lies in the plane of rotation of the doublet. (Right) Alignment of elongation major axis with the doublet axis. **d**. Schematics for interface shape decomposition into modes with different symmetries. **e**. Representative examples of interface shape for different rotation magnitudes, corresponding to points indicated in f. Top row: magnitude of rotational velocity and average interface deflection. Middle row: snapshots of E-cadherin labelled doublets with overlaid interface segmentation. Bottom row: Interface height map. **f**. Average interface deflection as a function of magnitude of rotational velocity. **g**. Average relative magnitude of the interface deformation modes. **h**. Correlation of orientation of deformation mode with direction of rotation (indicated on the left). Statistical tests: ***, *p* < 10^-4^. Error bars: 95% confidence interval of the mean. *n* = 14 doublets, *N* = 3 biological repeats.

## Mode decomposition of interface deformation

We then quantified the shape of the interface between the two cells of the doublet (Fig. 2d). We found that the average deviation of the interface from a planar shape was increasing with the magnitude of doublet rotation (Fig. 2e, f). Cross-sections of the doublet showed that the interface was curved in a way that evoked a yin-yang shape, as noticed previously for doublets rotating on a substrate^7^ (Fig. 1b–d). However, when looking at the full interface three-dimensional shape, we noticed that the shape was more complex than suggested by this simple picture (Fig. 2e). To make sense of this complex shape, we decomposed it into basic modes of deformations, which we obtained from Zernike polynomials and classified according to their symmetry properties (Fig. 2d, Supplementary Information section 4). We defined a “bowl” mode corresponding to a rotationally symmetric deviation of the interface; a “saddle-node” mode with the symmetry properties of a nematic, a “three-fold” mode with a three-fold rotation symmetry; and a “yin-yang” shape with a positive and a negative peak of deformation and the symmetry property of a vector (Ext. Fig. 4, Supplementary Information section 4.2). We then measured the average magnitude of these 4 deformation modes (Fig. 2g). We found that all 4 modes contributed to the interface shape and the saddle-node mode was dominating in relative magnitude (Fig. 2g). We then tested whether the orientations of the shape deformation modes were correlated with the axis of rotation (Fig. 2h). We calculated correlation values considering the symmetry properties of each mode of deformation. Only the yin-yang mode, but neither the saddle-node nor the three-fold mode, has an orientation correlated with the direction of rotation (Fig. 2h). Altogether the doublet interface has a complex three-dimensional shape, and one mode of interface deformation correlated with the doublet rotation.

## Cortical myosin distribution encoded in cell polarity

Having characterized doublet rotation and the doublet interface shape, we then asked whether key proteins of the cytoskeleton and adhesion machinery had a distribution correlated with the doublet rotation. We stained cell doublets for phosphorylated Myosin-Regulatory Light Chain (p-MRLC) and F-actin to label the acto-myosin cytoskeleton, and E-cadherin to label cell-cell contacts (Fig. 3a and Methods). Strikingly, we found that while E-cadherin and F-actin were largely concentrated at the cell-cell interface, p-MRLC was almost absent at the interface but concentrated at two bright zones near the boundary of the cell-cell junction. Immuno-fluorescence paxillin staining and live imaging of vasodilator-stimulated phosphoprotein (VASP) further revealed that focal contacts were also concentrated near the interface boundary (Fig. 3b, c). Live imaging combining markers for actin and myosin cytoskeleton, cell-cell adhesion, and focal contacts showed that myosin clusters and focal contacts appear distributed within the cortex on opposite sides near the doublet interface, judging from their position relative to the interface shape (Fig. 3d, e, Supplementary Video 5-7, Ext. Fig. 3, Supplementary Information section 6). Finally, the observed myosin dynamics (Ext. Fig. 3c, d, Supplementary Video 7) shows signs of cluster movements, with typical cluster velocities around 0.1-1 μm. min^-1^.

**Figure 3.**
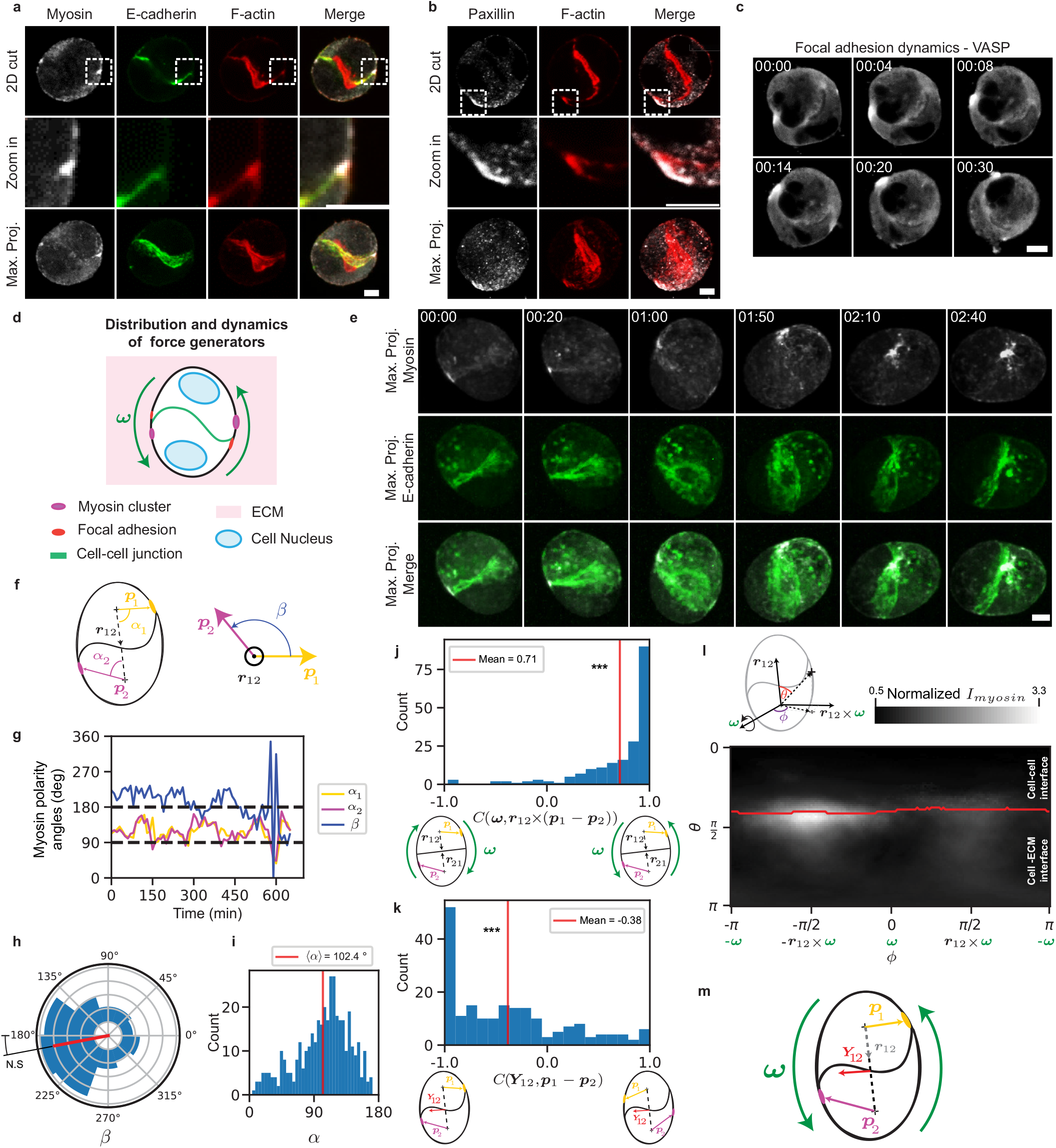
Distribution and dynamics of the force generators and adhesions in cell doublet. Snapshots of representative examples of distribution of (**a**) myosin, E-cadherin, F-actin and merge, and (**b**) paxillin, F-actin, and merge. *n* > 10 doublets for a and b. **c**. Dynamics of focal adhesion in the rotating doublet, labelled with VASP-GFP (grey). Time relative to the beginning of Movie 6. **d**. Schematics of distribution of force-generating and adhesion proteins in the rotating doublet. **e**. Snapshots of a rotating doublet with labelled E-cadherin-mNG (green) and myosin-KO1 (grey). **f.** Scheme of myosin polarity angles *α*_1_, *α*_2_ and *β*. **g**. Myosin polarity angles with respect to the doublet axis as a function of time for a single doublet. **h**. Histogram of *β*, the angle between the polarity vectors of cells 1,2, projected on the plane orthogonal to ***r***_12_. Red line, average orientation of the distribution. **i**. Histogram of *α*, the polarity angle relative to the doublet axis. **j**. Histogram for the correlation between the rotation vector ***ω*** and the cross-product between the doublet axis and the difference of cell polarities. Red line: average. **k**. Histogram for the correlation between the yin-yang orientation vector and polarity difference. Red line: average. **l**. Map of average myosin intensity in spherical coordinates, in a reference frame defined by the rotation vector ***ω***, the axis of the doublet ***r***_12_ and their cross-product. **m**. Summary of orientation of myosin polarities and yin-yang interface deformation mode. Scale bars: 5 μm. Time in hh:mm. Statistical tests: ***, *p* < 10^-4^. *N* = 3 biological repeats. Panel h-l: *n* = 12 doublets.

To quantify the cortical myosin distribution, we defined a polarity vector ***p*** associated with each cell, which was obtained from myosin signal intensity on the cell surface, corrected for a gradient in the z direction away from the microscope objective (Ext. Fig. 4c–e, Supplementary Information section 5). In line with our observation of myosin clusters, the two cell polarity vectors were consistently pointing away from the axis joining the doublet at an angle of nearly 90°, in opposite directions in each cell of the doublet (Fig. 3f–i). The cortical myosin polarity axis was correlated with both the doublet rotation axis and with the orientation of the yin-yang interface deformation (Fig. 3j–k). To better visualize the distribution of cortical myosin, we averaged the myosin intensity profile in a reference frame defined by the rotation vector and the cell doublet axis; this again revealed a strong myosin accumulation near the cell-cell interface, towards the direction of cell motion (Fig. 3l, Ext. Fig. 5). Overall, we conclude that the cortical myosin accumulates in clusters whose positions are correlated with rotation and interface deformation (Fig. 3l).

## An active surface model recapitulates doublet rotation

We then wondered if we could understand how doublets were physically rotating and the complex shape of their cell-cell interfaces (Fig. 4a and Ext. Fig. 6). We considered a physical model where each cell is described as an active viscous surface, subjected to active tension (Supplementary Information, section 7). Cell-cell adhesion is described by an interaction potential between cell surface points, with short-range repulsion and intermediate-range attraction. Surface flows are obtained from force balance equations at the cell surface, taking into account an external friction force proportional to the cell surface velocity. Polarity vectors were assigned to each cell, opposite to each other and oriented with a constant angle away from the cell-cell interface. A profile of active tension was imposed around the cell polarity axis (Supplementary Information, section 7.3). This profile was set using measurements of cortical myosin intensity in a reference frame defined by the cell polarity vector and the axis joining the doublet cells (Fig. 4b–c, Ext. Fig. 5, Ext. Fig. 6a–c, Supplementary Information section 5.4). Simulations were performed using the Interacting Active Surfaces (IAS) numerical framework^17^ (Fig. 4d, Supplementary Information, sections 7.1 and 7.2). Solving for the doublet dynamics, we found that a cortical flow emerges, the cell doublet rotates, and the cell-cell interface acquires a deformed shape (Fig. 4e, Ext. Fig. 6f and Supplementary Video 8). The simulated interface shape was a pure yin-yang deformation correlated with the direction of rotation (Fig. 4e–f), the doublet was elongated, and the major axis of elongation was rotating with the doublet (Fig. 4g), as experimentally observed (Fig. 2c, h). Other deformation modes were however absent, in contrast to experiments (Fig. 4e). Both the simulated rotational velocity and magnitude of interface deformation increase with the magnitude of the active tension deviation (Fig. 4h, i and Supplementary Information section 7.4). To test this prediction, we compared the magnitudes of the rotation and yin-yang interface deformation mode to the variation in cortical myosin polarity and found that they were indeed correlated (Fig. 4h, i). The simulated rotation magnitude was comparable to experiments for parameters giving rise to cortical flows of ~0.1μm/min, comparable to the observed speed of myosin clusters (Ext. Fig. 6f, Ext. Fig. 3d).

**Figure 4.**
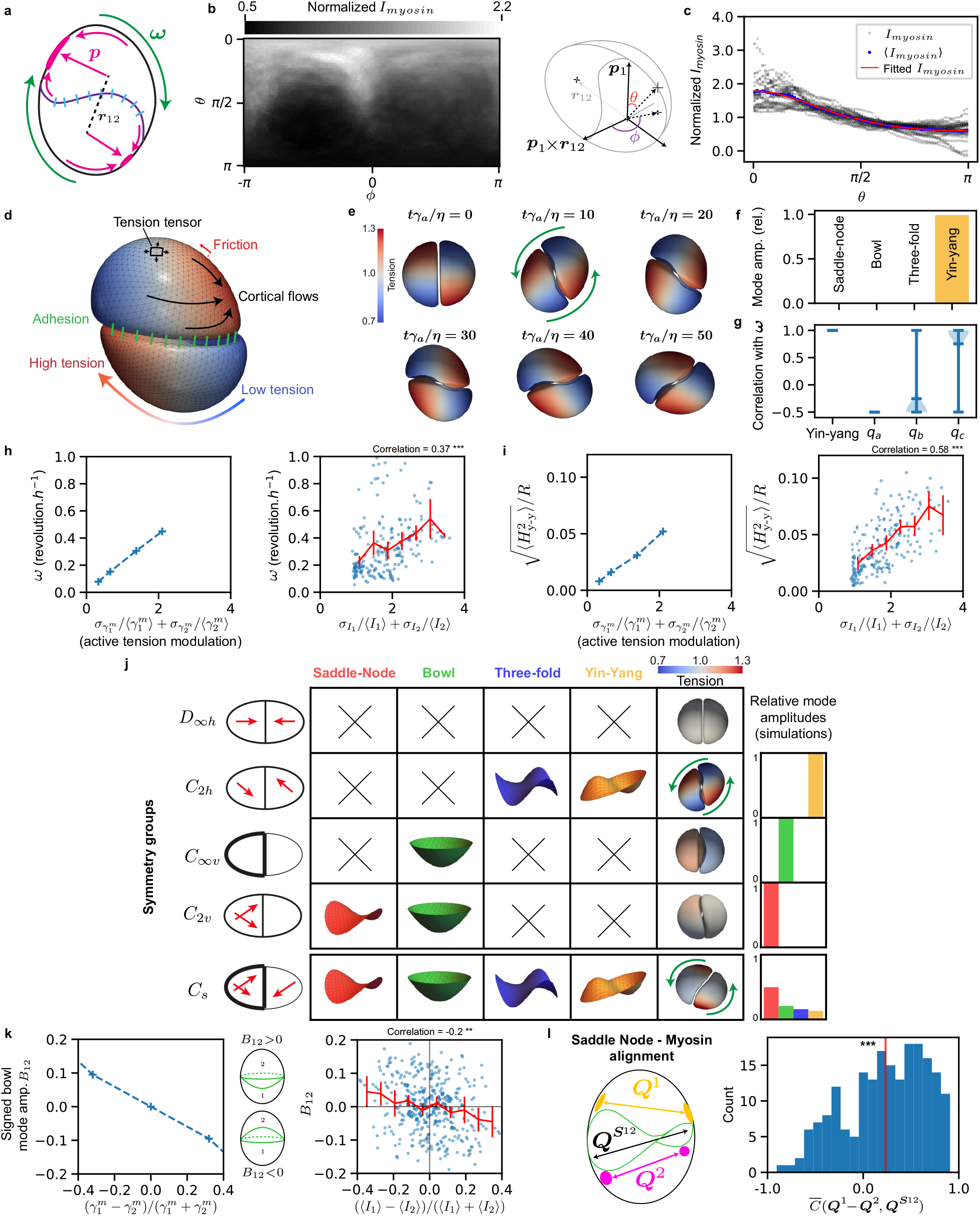
Interacting active surface simulations of a rotating doublet. **a**. Schematics of doublet rotation. **b**. Map of average experimental myosin intensity. **c**. Profiles of myosin intensity as a function of the angle *θ* defined in panel b. Grey: time average profile of individual cells, Blue: profile averaged over all cells, Red: fitted profile. **d**. Schematics for IAS simulation of a rotating doublet. **e**. IAS simulation results (*η* cortical viscosity, *γ*_a_ reference active tension). Time relative to the beginning of Movie 8. **f.** Amplitude of mode deformation for the simulated cell-cell interface in e. **g**. Correlation of yin-yang mode orientation, and doublet principal elongation, with direction of rotation. **h**. Left: rotation velocity of simulated doublet as a function of active tension modulation (*η*/*γ_a_* =1min). Right: Experimental rotation velocity as a function of myosin variation, Red: average rotational velocity. **i**. Left: Amplitude of yin-yang deformation mode as a function of the amplitude of the active tension modulation. Right: Experimental amplitude of yin-yang deformation mode as a function of myosin variation, Red: binned averages. **j**. IAS doublet simulation for cortical myosin profiles consistent with doublets in different symmetry groups (Schoenflies notation on the left). **k**. Left: Correlation of “bowl” mode amplitude with difference of average rescaled active tension. Right: Experimental correlation of bowl mode amplitude with difference of average myosin intensities. Red: binned average **l**. Experimental correlation of saddle-node mode with difference of cortical myosin nematic tensor. Error bars: 95% confidence interval of the mean. See Supplementary Information section 7. Statistical tests: **: *p* = 3.10^-4^, ***: *p* < 10^-4^.

## Curie principle applied to doublet interface shape

We then wondered how we could explain the emergence of modes of interface deformation other than the yin-yang shape. We reasoned that the Curie principle, stating that “the symmetries of the causes are to be found in the effects”^18,19^, implies that molecular cues guiding interface deformations should satisfy symmetry rules consistent with the observed interface shape (leaving aside the possibility of spontaneous symmetry breaking). We classified a set of configurations of doublets and polarity axis according to their symmetry properties (Fig. 4j). A configuration where cell polarities are in the same plane but shifted in opposite directions away from the doublet axis, belongs to the *C_2h_* point group in Schoenflies notation^20^. As a result, such a doublet should exhibit yin-yang and three-fold interface deformation, as observed in simulations (Fig. 4f, Ext. Fig. 6d). In contrast, tension asymmetry between the two cells of the doublets should give rise to the bowl deformation mode; while a nematic configuration of active tension distribution, with different intensities in each cell, should give rise to the bowl and saddle-node deformation mode (Fig. 4j). Simulating doublets with varying profiles of active tension confirmed these predictions (Fig. 4j). We then verified if this relationship between modes of cortical myosin distribution and modes of interface deformation could be observed in experiments. Indeed, we found that the magnitude of the bowl deformation mode was correlated to the difference in average cortical myosin intensity of the two doublet cells (Fig. 4k). We also noticed that the distribution of cortical myosin had a secondary, less concentrated cluster opposite to the main myosin cluster (Fig. 4b, Ext. Fig. 5). We reasoned that this secondary cluster was giving rise to a nematic distribution of cortical myosin, quantified by a nematic tensor^21^. Indeed, we measured a positive correlation between the nematic tensor of the saddle-node interface deformation mode, and the difference of cortical myosin nematic tensor between the two doublet cells (Fig. 4l). We then verified that simulating a doublet with an active tension profile summing a polar, nematic distribution and a difference in average tension between the two doublets cells, resulted in a complex interface shape with a similar mode decomposition as in experiments (Fig. 4j, last row, Fig. 2g). We conclude that the complex shape of the doublet interface can be understood, based on symmetry principles from the cortical myosin distribution in the doublet.

## Deletion or ectopic induction of myosin clusters affects doublet interface and motion in experiments and in simulations

We then reasoned that if the myosin clusters are responsible for cell rotation and interface deformation, perturbing their activity and localization would affect the cell doublet shape and motion. Indeed, treatment with the myosin inhibitor blebbistatin (Fig. 5a) resulted in rotation arrest and simultaneous flattening of the interface (Fig. 5a–c and Ext. Fig. 7). The effect of the blebbistatin-induced arrest of rotation was reversible: when we washed out the inhibitor, doublets retrieved rotational motion and a bent interface (Ext. Fig. 7 and Supplementary Video 9). We then aimed at specifically altering the two opposite myosin spots using laser ablation (Fig. 5d). This induced transient arrest of the rotation and interface flattening. Following a lag time of ~ 10 minutes, rotation restarted with a simultaneous increase in rotation velocity and interface deformation (Fig. 5e and Supplementary Video 10). Suppressing the gradient of active tension in simulation also suppressed rotation (Fig. 5d, e and Supplementary Video 11). In order to generate additional ectopic local myosin clusters, we then engineered a stable optogenetic cell line that we used to locally activate Rho (Fig. 5f and Supplementary Video 12, see Methods). Transient Rho activation resulted in ectopic myosin activation at the cell cortex comparable in intensity and size to spontaneous myosin clusters. This new cluster triggered the displacement of the doublet away from the region of activation while the doublet kept rotating (Fig. 5f–h). Introducing an ectopic region of increased active tension in simulations resulted in a similar drift of the doublet (Fig. 5f-h and Supplementary Video 13). Altogether, these results support the role of myosin clusters for driving doublet rotation.

**Figure 5.**
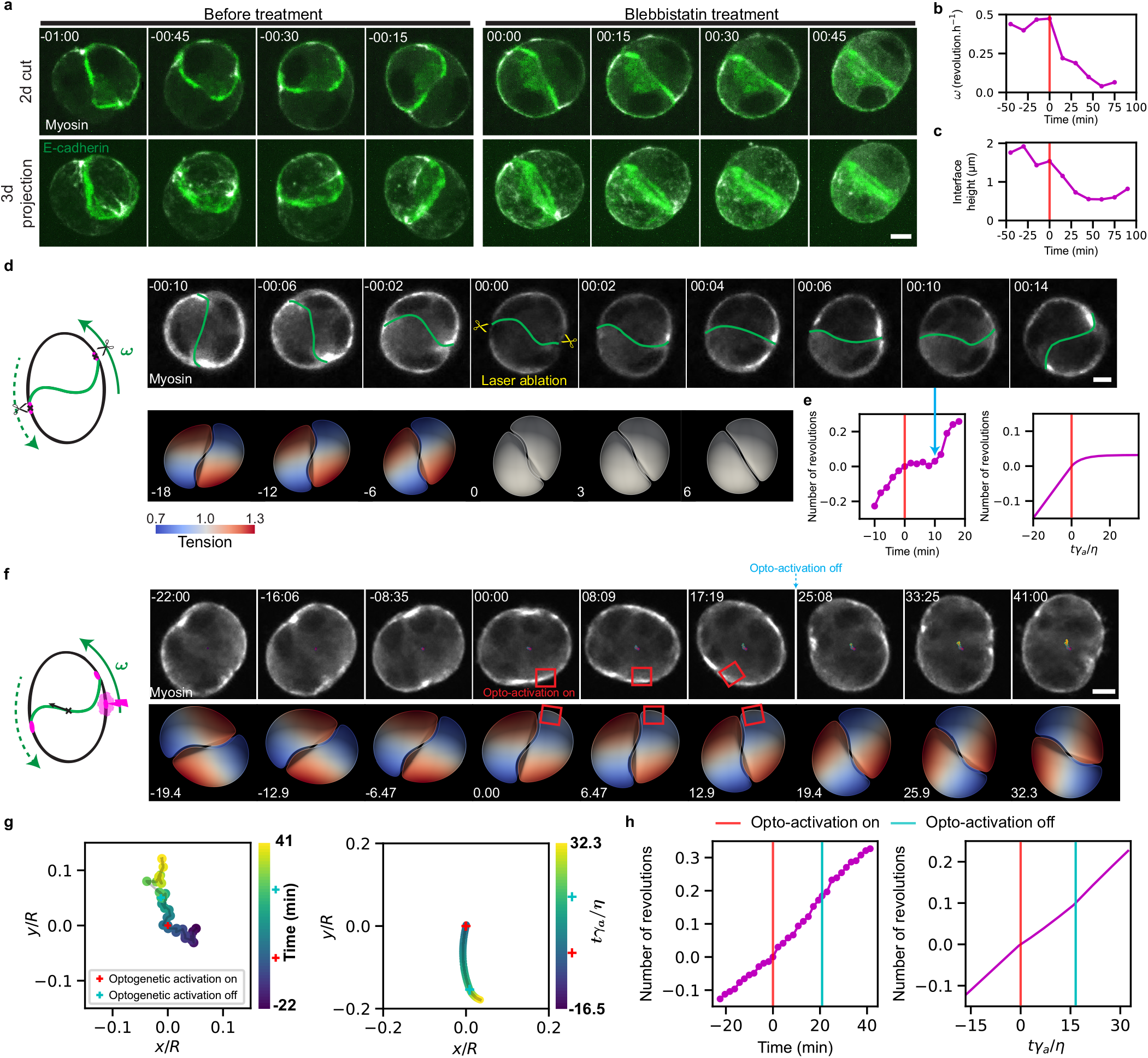
Perturbing the rotation motor. **a**. Snapshot of a rotating doublet, before and after Blebbistatin treatment. Green label: E-cadherin-mNG, grey label: MRLC-KO1 (see also Movie 9). *n* = 16 doublets. **b**. Magnitude of rotation as a function of time, before and after treatment with Blebbistatin. **c**. Magnitude of interface height as a function of time, before and after treatment with Blebbistatin. **d**. Snapshot of a rotating doublet labelled with MRLC-GFP (grey) (top: experiments, bottom: simulation), before and after laser ablation of myosin spots at time 0; ablation spots are indicated by two scissors (see also Movies 10 and 11). **e**. Number of visible turns as a function of time before and after laser ablation of myosin spots; time of ablation indicated by a red line and blue arrow shows the onset of rotation. **f**. Snapshot of a rotating doublet labelled with MRLC-iRFP (grey), before and after optogenetic activation of RhoA (red square) generating a local myosin cluster (top: experiments, bottom: simulation, time 0 activation) (see also Movies 12 and 13). **g**. Number of visible turns as a function of time, before, during and after optogenetic generation of a myosin cluster (*n* = 4 doublets) (left: experiments; right: simulations). **h**. Trajectory of the doublet’s center of mass before, during (red line) and after (blue line) optogenetic generation of a myosin cluster (experiment) and induction of a region of increased active tension in one cell (simulation). Scale bars: 5 *μ*m. Time in hh:mm, panel a,d. Time in mm:ss in panel f.

## Discussion

Our analysis shows that doublet rotation arises from myosin clusters positioned away from the axis joining the two doublet cells. Therefore, the doublet cell rotation requires cell-cell interactions to trigger the shift of the cell polarity axis. Consistent with this picture, in our experimental setup single cells do not rotate. It has been reported that in a bilayered Matrigel, which provides an external polarization axis, single MDCK cells rotate^11^. In this situation, the environment is providing a preferred direction. It would be interesting to track the cortical myosin distribution in cells rotating in these conditions to test if the polarity-based mechanism we propose also applies in that context.

Which mechanisms result in myosin cluster formation? Clusters could emerge by spontaneous symmetry breaking from an initially symmetric configuration where the cell polarities are pointing towards each other (Fig. 4j). The dynamics of increase of rotation magnitude after cell division (Fig. 1h), which resembles an exponential increase followed by saturation, is consistent with such a scenario. The position of myosin clusters could be related to protrusions which are sent by each cell beyond the interface at the cortex. Alternatively, they form in response to cell interface deformation, and induce rotation and further cell interface deformation, with the positive feedback loop at the origin of the instability and spontaneous symmetry breaking necessary for rotation. Among the deformation modes we have analyzed, the yin-yang mode has the right symmetry property to be coupled to a shift of the two cell polarities away from the doublet axis, which occurs in opposite directions in each cell of the doublet (Fig. 4j). Interestingly, we also observed that following cell division, a myosin-dense cluster forms at the center of the doublet interface and appears then to relocate towards the periphery of the contact (Ext. Fig. 8 and Supplementary Video 14). We note that myosin clusters have also been reported to generate stress in cytokinesis^22^. We also observe that focal contacts are polarized within each cell of the doublet, such that a myosin cluster in one cell is in close vicinity to a region dense in focal contacts within the opposite cell. This organization suggests either a common origin for myosin and focal contact distribution or a polarization mechanism relying on negative feedback between cortical myosin and focal contacts. Possibly, forces resulting from the cortical myosin inhomogeneous distribution promote focal contacts through a reinforcement mechanism^23,24^.

Our study shows that epithelial cell doublets allow to bridge the gap between microscopic players involved in cell motion and collective tissue dynamics. We propose that the spontaneous rotation we observed here is a manifestation of basic principles of cell interactions, involving cross-talks of cell polarity between neighbouring cells and polarity-oriented mechanical interactions between the cells and their environment. The ubiquity of collective rotations observed in various cell types *in vitro* suggests that they indeed emerge from generic principles. Analysis of the symmetries, here through the Curie principle, helps in making sense of these complex interactions. It would be interesting to see how basic rules of cell polarity interactions combine with mechanical forces to generate tissue self-organisation beyond collective cell rotation.

## Methods

### Cell culture and cell lines

Cells were maintained at 37°C in a 5% CO_2_ incubator. All following steps are performed under a sterile hood.

The maintenance of MDCK II cell lines was done using high Glutamax Modified Eagle’s Medium (Gibco, ref. 41090-028), 5% v/v Fetal Bovine Serum (South America Gibco, ref. 10270106), 1% v/v Non-Essential Amino Acid (Gibco, ref. 11140-050), 1% v/v sodium pyruvate (Gibco, ref. 11360-039) and 1% Penicillin-Streptomycin. MDCK cells were replated every 3 to 4 days when they reached 70-95% confluency. Trypsin-EDTA was used to detach the cells from the plate and with a seeding density of 5.10^4^ per cm^2^. To generate doublets, single cells were embedded in a Matrigel close to the surface. Coverslips were activated by O_2_ plasma and 100% Matrigel (Corning BV: 356231) was then used to coat coverslips. Following a 10 min incubation at 37°C, 100% Matrigel was polymerized and formed a basal layer. Single cells were deposited at a density of 15,000 cells/cm^2^. After 10 minutes incubation, unattached cells and excess medium were removed. Next a drop of 10 μl 100% Matrigel was deposited. After polymerization of Matrigel at 37°C in the incubator, culture medium was further added.

We prepared the following stable cell lines: MDCK II E-cadherin-GFP/Podocalyxin-mScarlett/Halo-CAAX to visualize the cell-cell junction and the lumen. To generate new myosin clusters by opto-genetics, we prepared a stable MDCK II cell line expressing iLID-LARG::mVenus^25^, engineered to be membrane-anchored by a slowly diffusing Stargazin membrane anchor^26^ combined with the DH/PH domain of the Leukemia-associated RhoGEF (LARG)^27^). This cell line allowed us to activate Rho locally and trigger the local recruitment of myosin. We visualized Rho activity with an active Rho sensor (2xrGBD-dTomato) and myosin localization with MRLC-iRFP703. We also used the following cell lines: MDCK II VASP-GFP for tracking focal contacts^28^, MDCK II MRLC-KO1/E-cadherin-mNG to follow myosin and cell-cell junctions^29^, MDCK II MRLC-GFP^30^, MDCK II E-cadherin-GFP and MDCK II E-cadherin-DsRed^31^. These cell lines allowed us to track specific correlations between rotation and adhesion or cytoskeleton protein localizations. E-cadherin was also used for segmentation purposes. To visualize F-actin live, we also used SiR-Actin (Tebu-Bio, 251SC001). Each experimental condition (biological repeat noted N) was reproduced at least 3 times on at least 10 doublets (written n in captions).

### Drugs treatments and immuno-fluorescence staining

To investigate the role of myosin, we used blebbistatin (Sigma, B0560) to inhibit myosin with the following steps. Timelapses of samples were acquired under the microscope and the medium was then changed by a medium containing 10 μM of blebbistatin. Following 1 hour incubation, samples were washed 3 times with fresh medium. Samples were further imaged to visualize the eventual re-initiation of rotation.

For immunostaining^9^, samples were washed with PBS and then fixed with 4% paraformaldehyde (PFA) diluted in PBS for 15 minutes. To permeabilize cells, cells were incubated with 0.5% Triton-X-100 for 15 minutes and then a blocking solution made of 1% Normal Goat Serum in PBS 1X was added overnight. The primary antibody was added directly to the blocking solution for two days at 4°C. Following 3 washing steps, samples were stained with the relevant secondary antibodies for 2 hours at room temperature. We used the following primary antibodies: Anti-E-cadherin (Abcam, Ab11512), Anti-Phospho-Myosin Light Chain 2 (Cell signaling technology, #3674), Anti-Paxillin (Abcam, Ab32084), and Alexa FluorTM Phalloidin 488 (Thermo Fisher, A12379) to visualize F-actin. Samples were washed three times in PBS and mounted on a home-made sample holder system for imaging and conservation.

### Microscopy

High throughput imaging was done using a spinning disk microscope with an inverted Leica spinning disk DMI8 equipped with an Orca Flash 2.0 camera (2048*2018 pixels with a size of 6.5 μm) using a 63x glycerol objective (NA = 1.3). The microscope was equipped with an incubation chamber to maintain the samples at 37°C, 5% CO_2_ and 85% humidity conditions.

To record the initiation of rotation, MDCK doublets were imaged 5 hours after cell seeding. 3D stacks were acquired with a z-step of 1 μm and x-y resolution of 0.206 or 0.103 μm, every 10 min up to 12 hours.

Confocal imaging of fixed samples was performed using the same setup. Laser power and digital gain settings were unchanged within a given session to ease quantitative comparison of expression levels among doublets.

We locally activated Rho with the scanning head of a confocal microscope (Leica SP8-UV) with a 458 nm laser each 10 s for 20 min, taking an image every minute. We could follow RhoA activity with a Rho binding domain sensor for active Rho (2xrGBD::dTomato)^32^ and myosin with the MRLC::iRFP703 probe.

### Image analysis

3D segmentation of cells was performed using a custom written ImageJ Macro involving LimeSeg plugin^33^ (Supplementary Information section 1). The center of mass, surfaces of cells were computed from the segmented 3D meshes using a written Python code. For the velocity measurements, rotational velocity of cells was computed using the cell-to-cell center of mass vector, velocity of cell 1 and velocity of cell 2 (Supplementary Information section 2).

Segmentation of the interface was performed using a customized Python code from the 3D meshes of each cell in a doublet. The 3D point cloud of the interface was then fitted to a polynomial function of degree 3. Characterisation of interface deformation and shape was then performed on the fitted surface (Supplementary Information section 4).

The fluorescence intensity signal of myosin was extracted from images and attributed to vertices of 3D meshes. The myosin distribution was then characterised by a polarity vector and a nematic tensor. We correlated these descriptors to the interface shape and the doublet rotation (Supplementary Information section 5).

Cylindrical and planar projections of the basal (E-cadherin) and interfacial (myosin) signals were created using a custom Python code (Supplementary Information section 6).

For the visualization and 3D rendering, we used Paraview^34^ software.

### IAS simulation framework

Simulations of rotating doublets were performed using the IAS framework. A polarity is introduced for each cell which modulates the active tension profile along its surface. This leads to the emergence of a cortical flow which propels the cells through friction with the external medium. The integrity of the doublet is maintained using an adhesion interaction potential. See Supplementary Information section 7 for the complete description of the simulations and parameters used.

### Statistical tests

We used a bootstrapping approach for all the statistical tests performed. The tests in Figs. 2c,2h, 3h, 3j, 3k, 4h, 4i, 4l, were performed by generating 50 000 samples of mean using bootstrapping. For the correlation tests in Figs. 4h,i,k, we generated independent samples of the correlation defined as < (*a*-< *a* >) · (*b*-< *b* >) >/(*σ_a_σ_b_*) (for two signals *a* and *b* with *σ_a_,σ_b_* their respective standard deviations). This correlation takes values between −1 and 1. For Figs. 4k we used 200 000 samples. All the p-values correspond to one tailed tests.

## Supporting information

SV1_All aggregates rotate spontaneously

SV2_All doublets rotate with similar velocity

SV3_Doublet also rotate when two cells meet

SV4_A typical segmentation of cells

SV5_F-actin

SV6_Focal adhesion_Vasp

SV7_Myosin clusters localize near the cell-cell interface

SV8_Reference_simulation

SV9_Myosin activity is needed for rotation

SV10_Myosin clusters ablation

SV11_laser_ablation_simulation

SV12_Optogenetic_Activation_translates_doublet

SV13_optogenetic_simulation

SV14_Myosin redistribution after cytokinesis

## Acknowledgments

We thank Erwan Grandgirard and the Imaging Platform of IGBMC, and the Salbreux and Riveline groups for help and discussions. Q.V. and G.S. are supported by the University of Geneva. L.L. and T.G. are supported by HFSP and by the University of Strasbourg and by la Fondation pour la Recherche Médicale. D.R. acknowledges the Interdisciplinary Thematic Institute IMCBio, part of the ITI 2021-2028 program of the University of Strasbourg, CNRS and Inserm, which was supported by IdEx Unistra (ANR-10-IDEX-0002), and by SFRI-STRAT’US project (ANR 20-SFRI-0012) and EUR IMCBio (ANR-17-EURE-0023) under the framework of the French Investments for the Future Program. O.P. and D.R. thank funding from SNF Sinergia. D.R. and A.H. acknowledge the Research Grant from Human Frontier Science Program.

## Author contributions

G.S. and D.R. supervised the study. L.L., T.G. and D.R. conceived and analyzed experiments. L.L. performed the experiments with support from T.G., R.B., M.L.; O.P., K.U., and C.M.-L., A.H. contributed new optogenetic tool and cell lines respectively. T.G., Q.V. and G.S. performed data analysis. Q.V. and G.S. designed the theoretical model. Simulations were performed by Q.V. with support from A.T.-S. The manuscript was written by G.S. and D.R. based on joint discussions with L.L., T.G., Q.V.

---

**Ext. Fig. 1.**
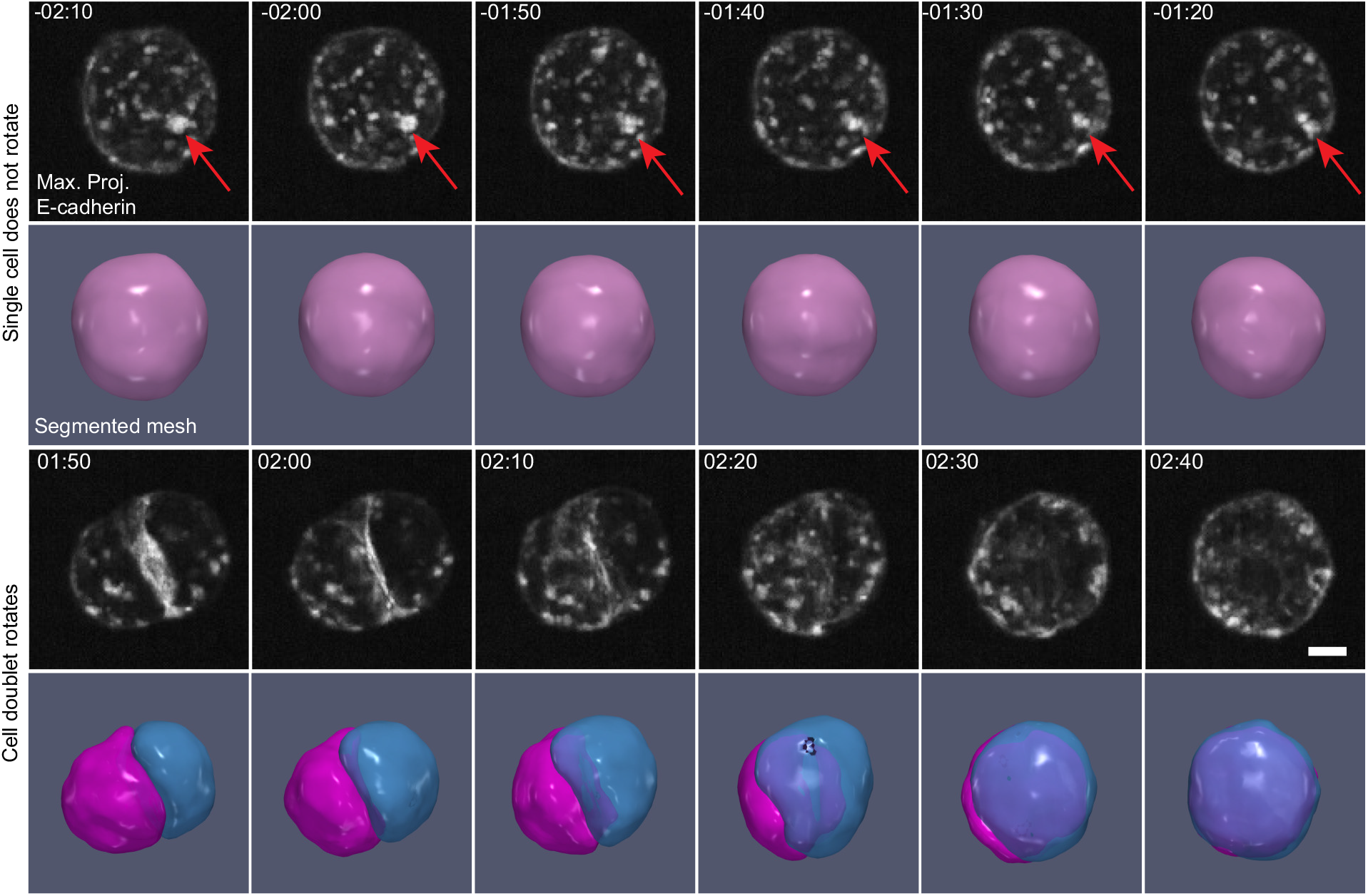
Single cells do not rotate. Snapshot of single cell (two top rows) and cell doublet (two bottom rows). Cells labelled with E-cadherin-mNG (grey) - see beginning of Movie 2. For each case, top row: maximum projection of E-cadherin, bottom row: cell segmentation. Time relative to cell division. Scale bars: 5 *μ*m. Time in hh:mm.

**Ext. Fig. 2.**
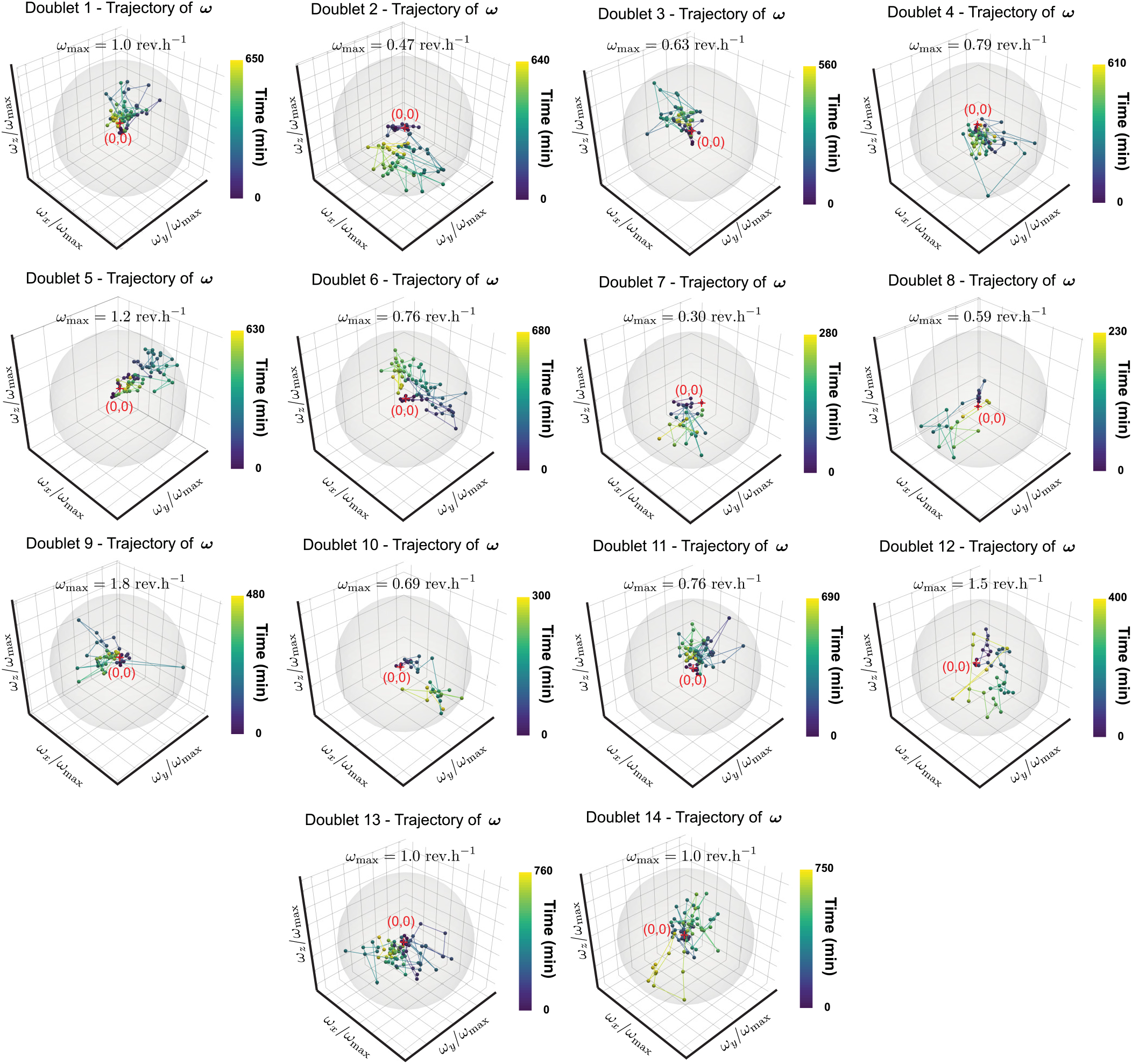
Trajectories of rotational velocities. Similar to Fig. 1h: trajectories of the rotation vector of cell doublets after cell division for all 14 doublets, normalized with respect to their respective largest amplitudes (corresponding to Movie 2). Grey sphere has unit radius.

**Ext. Fig. 3.**
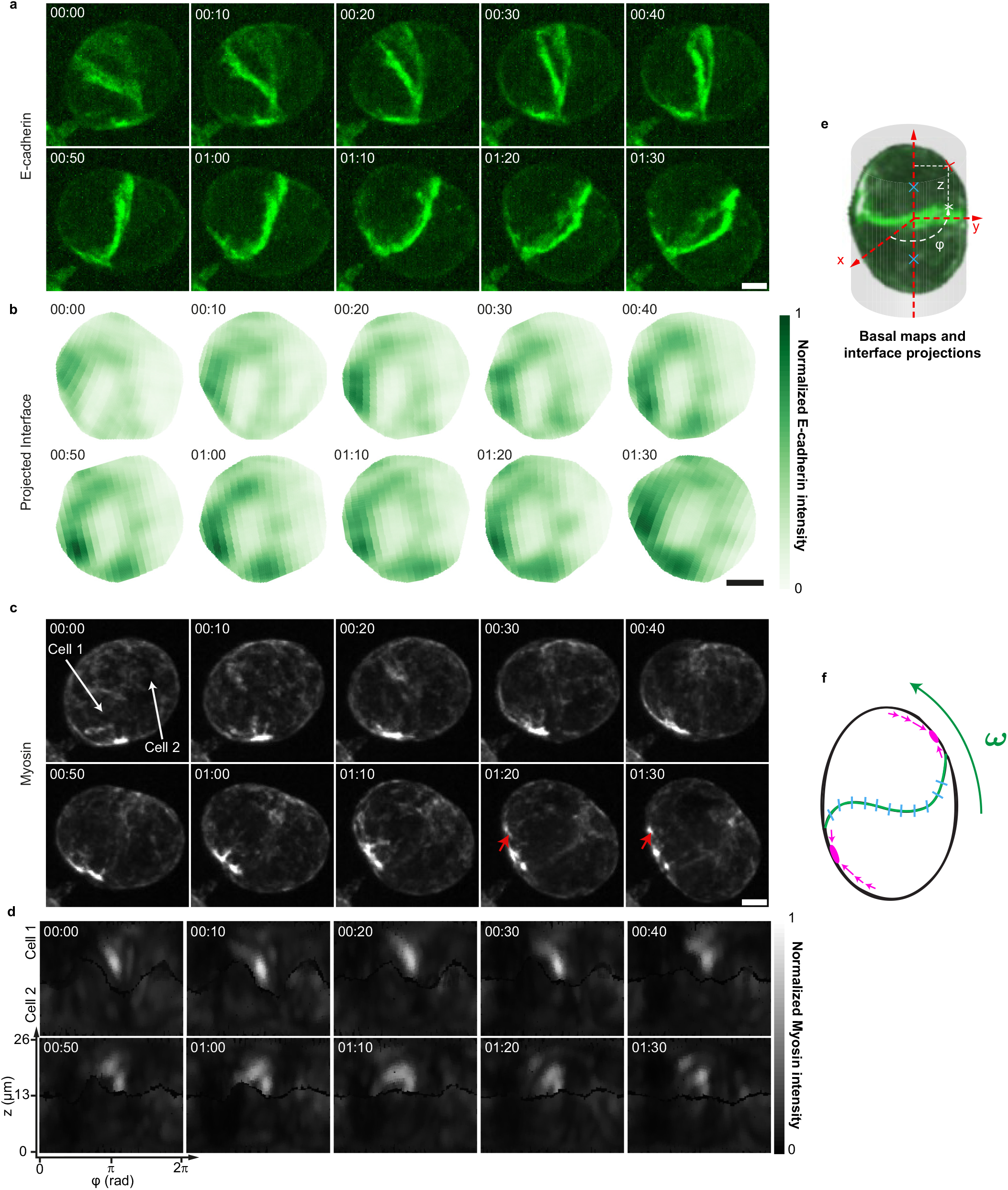
Patterns of E-cadherin at the cell-cell interface and myosin dynamics at the cortex. **a**. E-cadherin-mNG (grey) labelled rotating doublet. **b**. Patterns of E-cadherin distribution on the doublet cell-cell interface, viewed en-face, for the cell shown in a. **c-d**. Mapping of the myosin dynamics at the cortex (normalization described in SI section 6). Myosin clusters highlighted with red arrows exhibit a motion with a velocity of about 0.1*μ*m/min. (see Movie 7) **e**. Schematic for projection method. **f.** Schematics for myosin (purple) and cadherin (blue) distribution. Scale bars: 5 m. Time in hh:mm.

**Ext. Fig. 4.:**
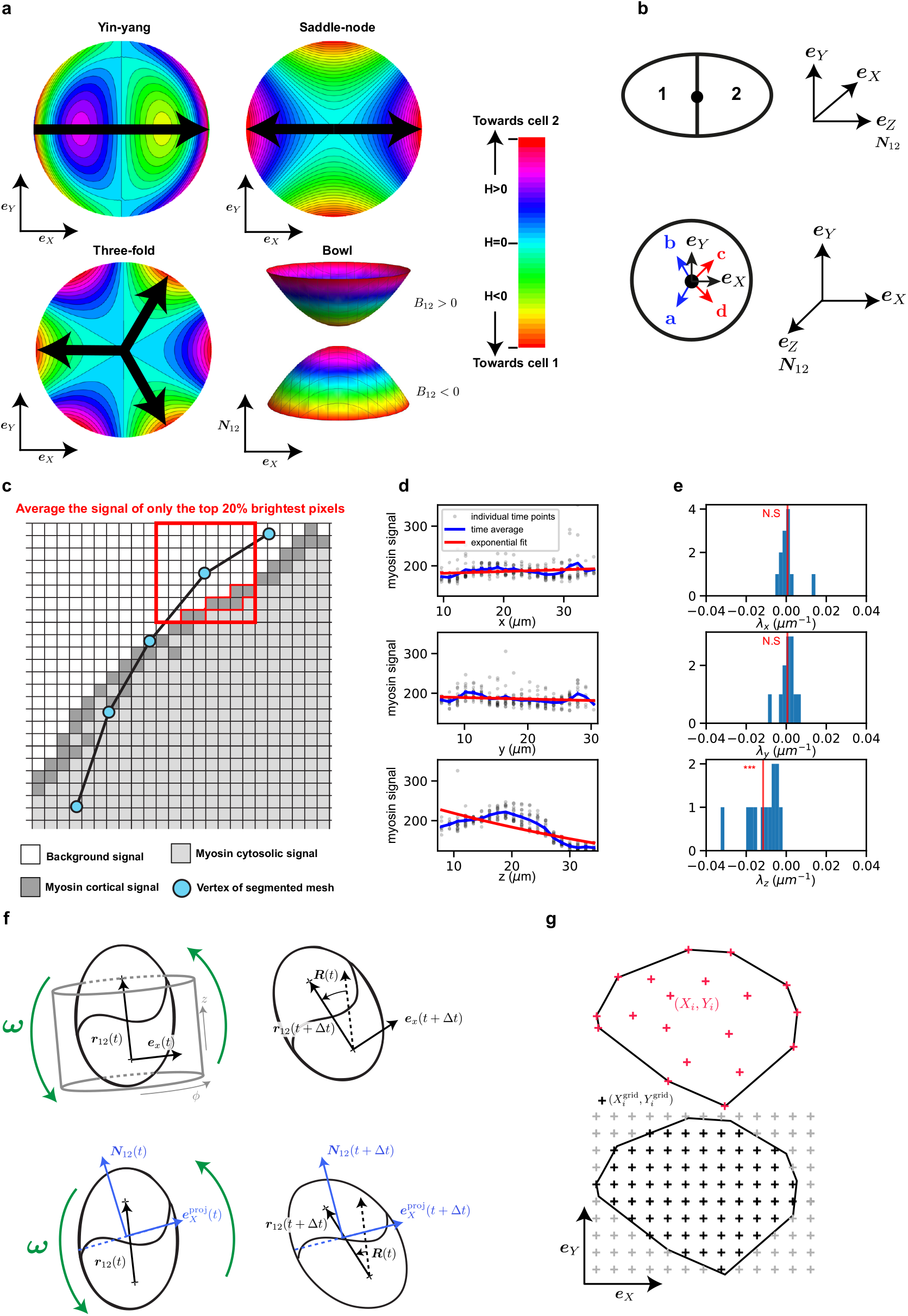
Analysis of interface shape. **a**. Height profile of interface deformation modes. The yin-yang orientation is characterized by a vector, the saddle-node by a nematic, and the three-fold by a three-fold orientational order. **b**. Schematic for the orientation of the vectors (***e**_X_, **e**_Y_, **e**_Z_ = **N***_12_) associated to the interface of the cell doublet. The vectors ***a**, **b**, **c**, **d*** are introduced to define transformations in Supplementary Table 1. **c**. Schematics of method used to obtain cortical intensities from cell segmentation (see Supplementary Information section 5.1) **d**. Average profile of myosin fluorescence intensity in the x, y, z directions, for a representative doublet. **e**. Histogram of fitting parameters characterizing the average myosin profiles, as in d, for all doublets. **f, g**. Procedure to create interfacial and basal maps. **f**. (Top) The vector ***r***12 is used as an axis around which to project the myosin signal intensity. The reference frame (***e**_x_, **e**_y_, **e**_z_*) defining the cylindrical coordinates (*z, φ*) rotates in a way that is consistent with the doublet rotation. (Bottom) A similar process is used for the interfacial maps of E-cadherin signal. A reference vector is rotated with the doublet to define a consistent viewpoint and is projected at each time *t* in the plane of the interface defined by ***N***_12_. **g**. (Top) 2D coordinates (*X_i_,Y_i_*) of the interface vertices i, surrounded by their convex hull. (Bottom) A regular grid of new coordinates 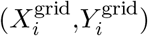 is created inside the convex hull (black points). ***, *p* < 10^-4^.

**Ext. Fig. 5:**
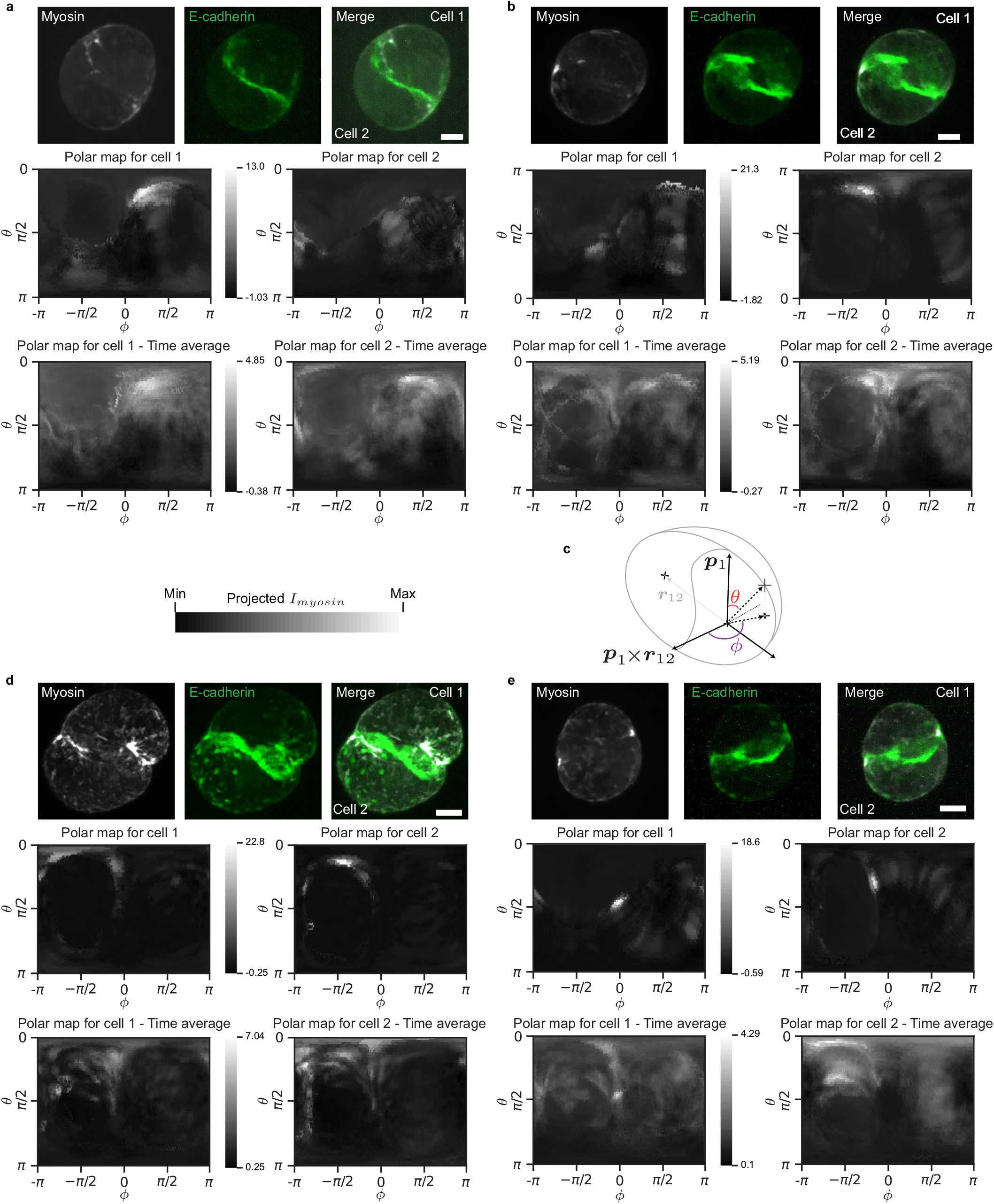
Example of polar maps of cortical myosin intensity. **a,b,d,e**. Maps of experimental myosin intensity after calibration in spherical coordinates, in a reference frame defined by the polarity axis ***p***, the axis of the doublet ***r***_12_ and their cross-product. For each example, top row: snapshot of doublet, maximum projections of myosin (MRLC-GFP), E-cadherin (Ecadherin-mNG) and merge, middle row: individual cell maps corresponding to the above snapshot, bottom row: time average cell maps corresponding to the time series from which the snapshot in the top row was taken from. **c**. Scheme of the reference frame. Scale bar: 5 *μ*m. Normalization described in Supplementary Information section 6.

**Ext. Fig. 6.**
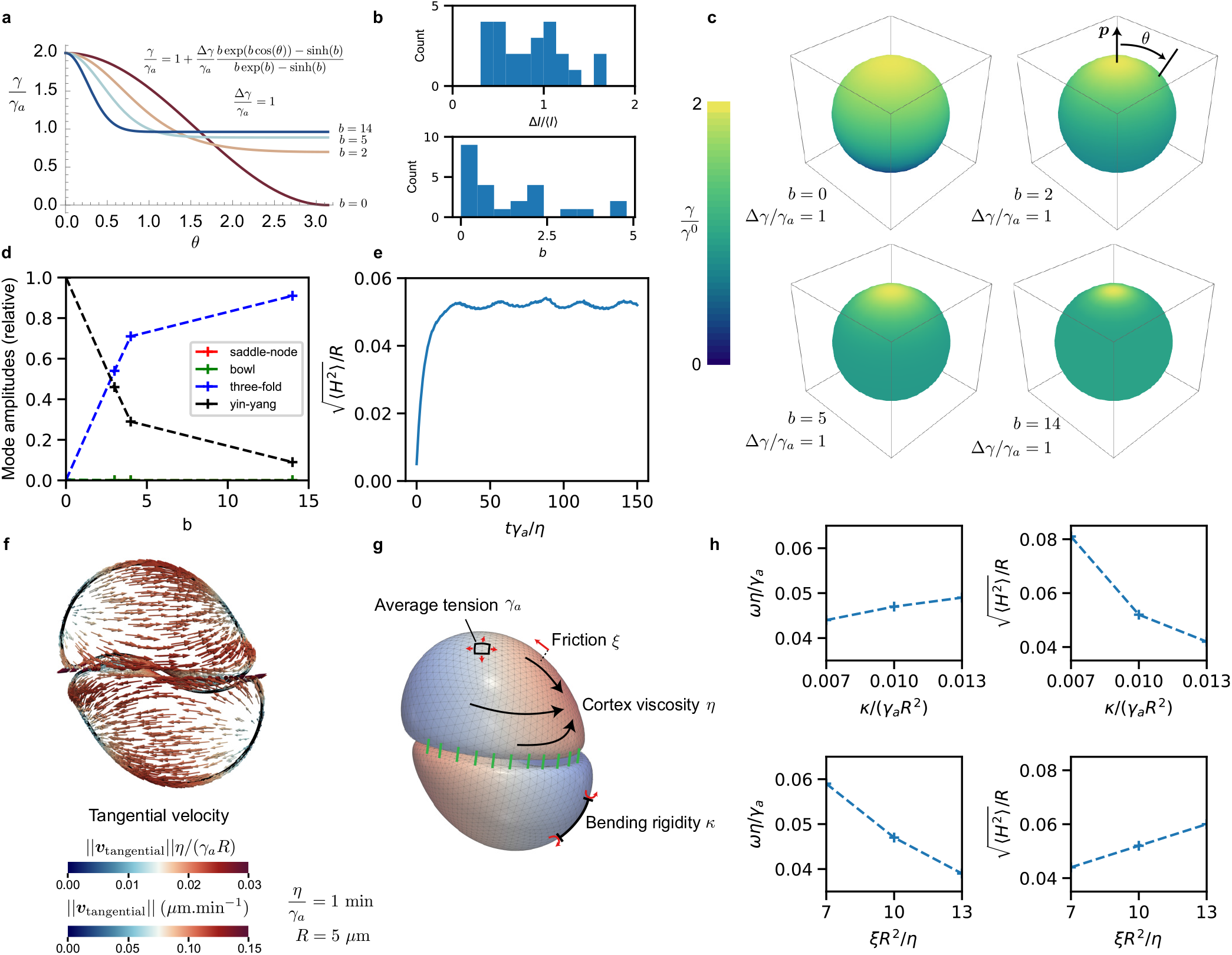
Additional IAS simulation results. **a**. Examples of tension profiles around the polarity vector, for different values of the parameter b, which controls the spread of the tension around the maximum value (see Supplementary Information section 7.3). **b**. Histogram of fitted values of *b* and Δ*I*/〈*I*〉 to temporally average myosin profiles for individual doublets, showing the distribution of myosin intensity magnitude and spot sizes. The fitting procedure and parameters are described in Supplementary Information section 7.6. **c**. Tension profiles displayed on spheres with values of b identical to panel a. Larger values of b correspond to smaller spots. **d**. Relative amplitudes of the deformation modes as a function of a varying active tension profile whose spread is determined by b. As the active tension spot size becomes smaller (larger values of b), the yin-yang mode is replaced by the three-fold mode. **e**. Interface deflection as a function of dimensionless time for the simulation shown in Fig. 4e. The interface deflection relaxes to a steady-state showing a slightly oscillatory behaviour. **f**. Cortical flow profile at steady-state for the simulation shown in Fig.4e. For *η*/*γ_a_*=1 min and *R* = 5*μ*m, the typical flow magnitude is ~ 0.1*μ*m.min^-1^. **g**. Explanatory scheme of IAS simulation, with key simulation parameters. **h**. Effect of varying the normalized friction coefficient *ξR^2^/η* and the normalized bending rigidity *κ*/(*γ_a_R*^2^) on the rotation velocity and the interface deflection, around parameter values chosen in simulations of Fig. 4, 5 and other panels of Ext. Fig. 7. See Supplementary Information section 7.5 for additional simulation parameters.

**Ext. Fig. 7.**
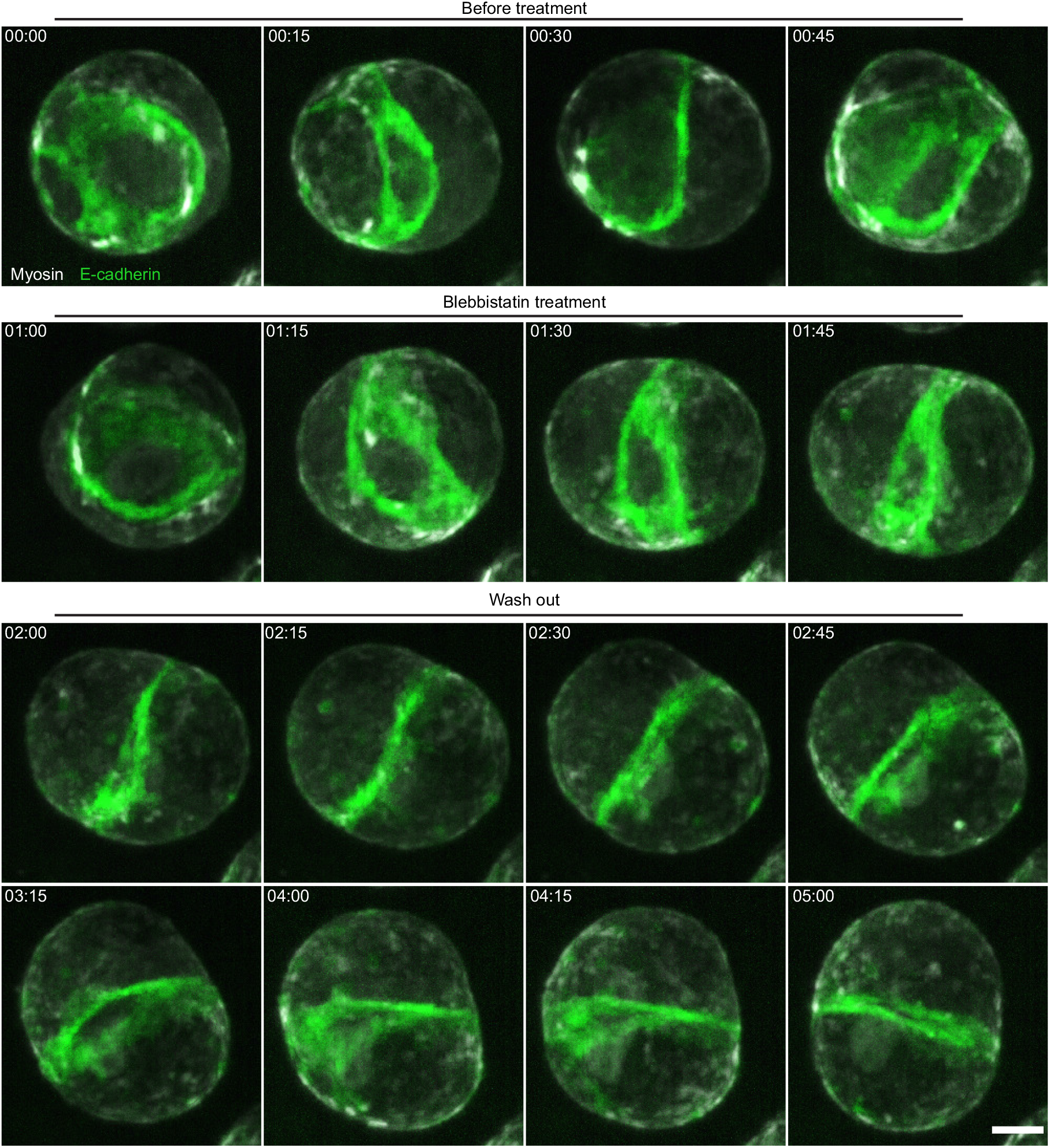
Blebbistatin treatment. Snapshots of blebbistatin experiment including before and after blebbistatin treatment followed by washout (see Movie 9). E-cadherin (green). Myosin (grey). Scale bar: 5 *μ*m. Time in hh:mm.

**Ext. Fig. 8.**
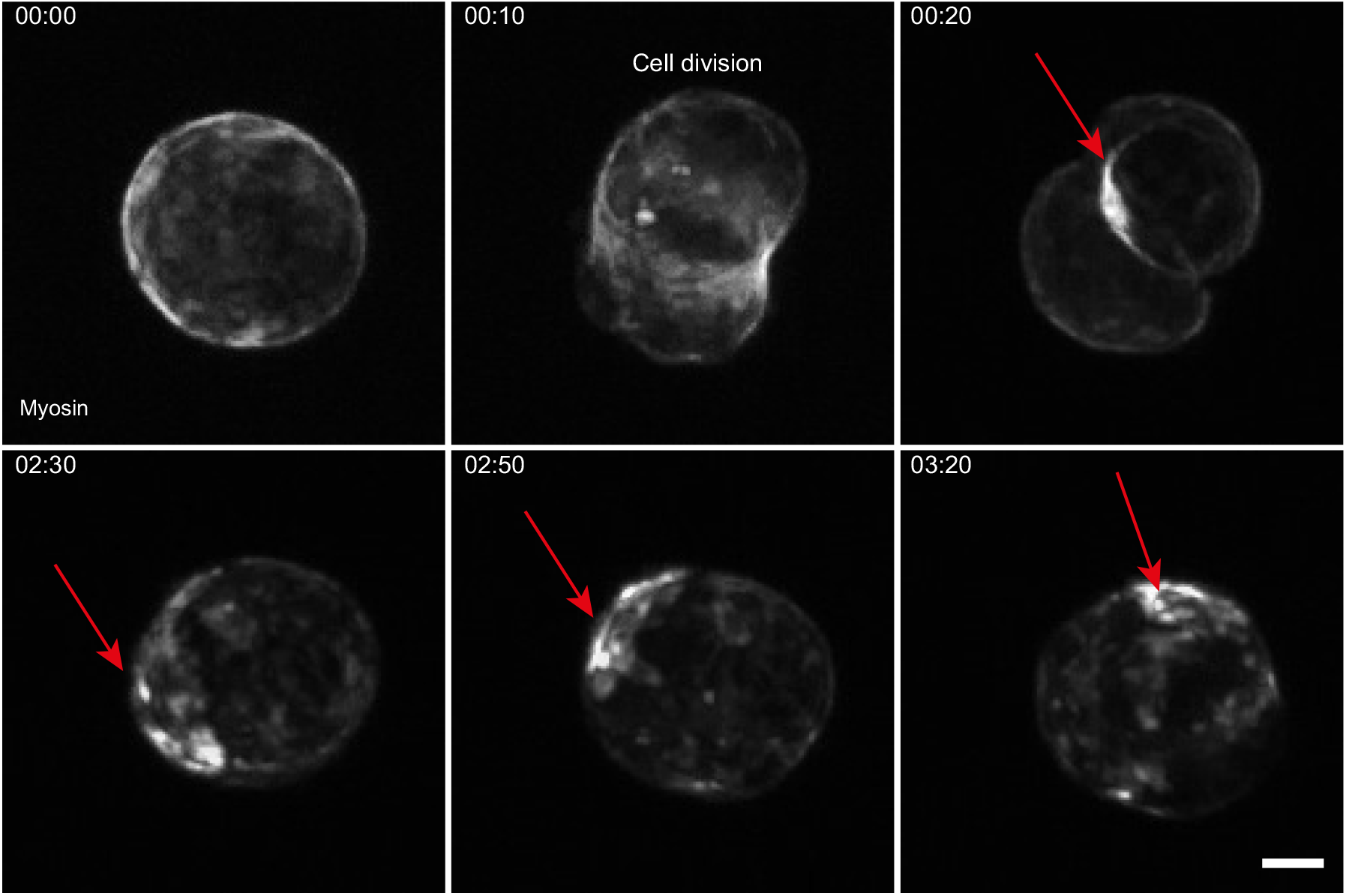
Myosin redistribution after cytokinesis. Snapshot of asymmetric myosin distribution before and after cell division (see Movie 14). Scale bar: 5 *μ*m. Time in hh:mm.

## Supplementary Videos legends

- Supplementary Video 1 - All doublets rotate spontaneously. MDCK cells expressing E-cadherin-mNG (in green) and Podocalyxin-mScarlett (in red). Time in hh:mm, scale bar: 20 *μ*m.
- Supplementary Video 2 - All doublets rotate with similar velocity. MDCK cells expressing E-cadherin-mNG (in grey). Time in hh:mm, scale bar: 5 *μ*m.
- Supplementary Video 3 - The doublet also rotates when two cells meet. MDCK cell expressing E-cadherin-GFP (in green) and E-Cadherin-DsRed (in magenta). Time in hh:mm, scale bar: 5 *μ*m.
- Supplementary Video 4 - A typical segmentation of cells. The doublet (left) expressing E-cadherin-mNG is shown next to its segmented version (right). Time in hh:mm, scale bar: 5 *μ*m.
- Supplementary Video 5 - F-actin localises at the cell-cell interface and within protrusions. MDCK cells expressing E-cadherin-mNG (in green) and F-actin labeled with SiR-actin (in grey). Time in hh:mm, scale bar: 5 *μ*m.
- Supplementary Video 6 – Focal adhesions localize near the cell-cell interface. MDCK cells expressing VASP-GFP (in grey). Time in hh:mm, scale bar: 5 *μ*m.
- Supplementary Video 7 - Myosin clusters localize near the cell-cell interface. MDCK cells expressing E-cadherin-mNG (in green) and MRLC-KO1 (in grey). Time in hh:mm, scale bar: 5 *μ*m.
- Supplementary Video 8 – Reference simulation shown in Fig. 4e. Cross-section of a rotating doublet showing a yin-yang interface deformation mode. The colormap indicates the active tension *γ/γ_a_* on the membranes. Dimensionless time *tγ_a_/η* is indicated. See Supplementary Information section 7.5 for simulation parameters.
- Supplementary Video 9 - Myosin activity is needed for rotation. The doublet rotates and stops its motion when blebbistatin is added (time 01:00); rotation starts again after washout. MDCK cells expressing E-cadherin-mNG in green and MRLC-KO1 in grey. Time in hh:mm, scale bar: 5 *μ*m.
- Supplementary Video 10 - Myosin clusters ablation corresponds to rotation arrests and changes in interface shape. MDCK cells expressing MRLC-GFP (in grey). Time in hh:mm, scale bar: 5 *μ*m.
- Supplementary Video 11 – Laser ablation simulation. Cross-section of a rotating doublet at steady-state, whose tension modulation is switched off at t=0. The colormap indicates the active tension *γ/γ_a_* on the membranes. Dimensionless time *tγ_a_/η* is indicated. See Supplementary Information section 7.5 for simulation parameters.
- Supplementary Video 12 - Local activation of Rho at time 0 (red square) leads to the generation of myosin clusters and this shifts the rotation to translation. The center of mass is tracked throughout the movie and is indicated with changing colours. MDCK cells expressing MRLC-iRFP. Time in hh:mm, scale bar: 5 *μ*m.
- Supplementary Video 13 – Optogenetic simulation. Cross-section of a simulated rotating doublet initially at steady-state. Active tension is increased in a spot in one of the cells, from time *tγ_a_/η* = 0 to *tγ_a_/η*=16.57. The added spot impairs the rotation, makes the doublet asymmetric and creates a drift of its center of mass. See Supplementary Information section 7.5 for simulation parameters.
- Supplementary Video 14 – Myosin clusters appear as a remnant from the cytokinetic ring. MDCK cells expressing MRLC-GFP (in grey). Time in hh:mm, scale bar: 5 *μ*m.

