## Supplementary Information for "Polarity-driven three-dimensional spontaneous rotation of a cell doublet"

December 20, 2022

#### Contents

**1 Segmentation of cell doublets** **2**

**2 Rotation vector computation** **2**

**3 Doublet shape analysis** **3**

**4 Interface shape analysis** **3**

### 1 Segmentation of cell doublets

Individual 3D cell segmentation was performed using a semi-automated segmentation method. The segmentation was done on Z-stack images ($\sim 40$ slices of thickness $\Delta z = 1.0$ $\mu$m, and in-plane resolution of $\Delta\ell = 0.206$ $\mu$m or 0.103 $\mu$m) of cells expressing fluorescent E-cadherin (E-cadherin mn-Green) that were obtained using spinning disk confocal microscopy. The 3D volumes were segmented using an ImageJ 3D image segmentation tool LimeSeg [1] with parameters that had to be adapted depending on the quality of the E-cadherin intensity signal. Prior to the segmentation computation, contour seeds were drawn on each z-slice to help the algorithm to converge. After each 3D analysis converged, each segmentation result was checked and confirmed by controlling how the boundary of the segmented cells fitted the original signal. In the case of cells expressing Myosin Regulatory Light Chain (MRLC) - MRLC-KO1 - as well as E-cadherin-mnG, both signals were used to check the quality of the segmentation. If the segmentation result was not accurate, a custom-made ImageJ macro was designed to enable the operator to re-draw the contour seeds or either vary the parameters used in LimeSeg. The coordinates of the vertices in the final meshes are given in dimensionless units $(x/\Delta\ell, y/\Delta\ell, z/\Delta\ell)$ with $(x, y, z)$ the coordinates of the vertices in $\mu$m.

### 2 Rotation vector computation

In general, one can decompose the movement of two points (of positions $\boldsymbol{r}_1(t)$ and $\boldsymbol{r}_2(t)$) into a translation with velocity $\boldsymbol{v}_t$, a rotation characterized by $\boldsymbol{\omega}$ and an extension with velocity $v_e$:

$$\begin{aligned} \boldsymbol{v}_1 &= \boldsymbol{v}_t - \frac{1}{2}\boldsymbol{\omega} \times \boldsymbol{r}_{12} - \frac{\boldsymbol{r}_{12}}{||\boldsymbol{r}_{12}||} v_e, \\ \boldsymbol{v}_2 &= \boldsymbol{v}_t + \frac{1}{2}\boldsymbol{\omega} \times \boldsymbol{r}_{12} + \frac{\boldsymbol{r}_{12}}{||\boldsymbol{r}_{12}||} v_e, \end{aligned} \tag{1}$$

with $\boldsymbol{r}_{12} = \boldsymbol{r}_2 - \boldsymbol{r}_1$, and $\boldsymbol{\omega} \cdot \boldsymbol{r}_{12} = 0$. The three components are uniquely defined as:

$$\boldsymbol{v}_t = \frac{\boldsymbol{v}_1 + \boldsymbol{v}_2}{2}, \; v_e = \frac{(\boldsymbol{v}_2 - \boldsymbol{v}_1) \cdot \boldsymbol{r}_{12}}{2||\boldsymbol{r}_{12}||}, \; \boldsymbol{\omega} = \frac{\boldsymbol{r}_{12} \times (\boldsymbol{v}_2 - \boldsymbol{v}_1)}{||\boldsymbol{r}_{12}||^2}. \tag{2}$$

The orientation of $\boldsymbol{\omega}$ corresponds to the axis of rotation (which is by definition perpendicular to $\boldsymbol{r}_{12}$), and its amplitude to the angular velocity. Since, experimentally, we only have access to discrete time points, we use a slightly different formula to estimate $\boldsymbol{\omega}$ from the cell displacements between time $t$ and $t + \Delta t$:

$$\boldsymbol{\omega}^{\Delta t} = \frac{1}{\Delta t} \frac{\boldsymbol{u}_{12}(t) \times \boldsymbol{u}_{12}(t + \Delta t)}{||\boldsymbol{u}_{12}(t) \times \boldsymbol{u}_{12}(t + \Delta t)||} \arccos(\boldsymbol{u}_{12}(t) \cdot \boldsymbol{u}_{12}(t + \Delta t)) \text{ with } \boldsymbol{u}_{12}(t) = \frac{\boldsymbol{r}_{12}(t)}{||\boldsymbol{r}_{12}(t)||}. \tag{3}$$

This formula is obtained by first establishing a differential equation on $\boldsymbol{r}_{12}$ derived from Eq. 1:

$$\frac{d\boldsymbol{r}_{12}}{dt} = \boldsymbol{\omega} \times \boldsymbol{r}_{12} + \frac{2\boldsymbol{r}_{12}}{||\boldsymbol{r}_{12}||} v_e. \tag{4}$$

This equation then translates into an equation for $\boldsymbol{u}_{12}(t)$:

$$\frac{d\boldsymbol{u}_{12}}{dt} = \boldsymbol{\omega} \times \boldsymbol{u}_{12}. \tag{5}$$

Assuming that $\boldsymbol{\omega}$ is constant between $t$ and $t + \Delta t$, and that $\boldsymbol{\omega} \cdot \boldsymbol{u}_{12}(t) = 0$, one can obtain Eq.3 by solving Eq. 5 between $t$ and $t + \Delta t$.

### 3 Doublet shape analysis

We quantify the elongation of a cell doublet as follows. We use segmentation meshes of the two cells of the doublet. The segmentation meshes are given by the vertices $\{\boldsymbol{v}_{1i} = \{v_{1ix}, v_{1iy}, v_{1iz}\}\}, \{\boldsymbol{v}_{2i}\}$ representing the surfaces of cell 1 and 2 of the cell doublet. We define a distance threshold of $d_{th} = 10$ in the dimensionless units defined in section 1 and select all vertices (from both cells) that are less than a distance $d_{th}$ away from the other cell. These vertices $\{\boldsymbol{v}_{\text{int}}\}$ are defined to be the interface vertices and will be used for the interface analysis in the next section. The remaining vertices $\{\boldsymbol{v}_{\text{outer},i} = \{v_{i,x}, v_{i,y}, v_{i,z}\}$ for $i \in \{1, ..., N_{\text{outer}}\}\}$ are used to define a shape tensor of the doublet:

$$\begin{aligned} Q^{3d}_{\alpha\beta} &= \frac{1}{N_{\text{outer}}} \sum_{i=1}^{N_{\text{outer}}} \left(v_{\text{outer},i,\alpha} - \langle v_{\text{outer},\alpha}\rangle\right)\left(v_{\text{outer},i,\beta} - \langle v_{\text{outer},\beta}\rangle\right) \text{ with } \alpha, \beta \text{ in } \{x, y, z\}, \\ \text{with } \langle v_{\text{outer},\alpha}\rangle &= \frac{1}{N_{\text{outer}}} \sum_{i=1}^{N_{\text{outer}}} v_{\text{outer},i,\alpha} \ . \end{aligned} \tag{6}$$

Diagonalising the tensor provides three eigenvalues $w_a \geq w_b \geq w_c$ associated with three orthonormal eigenvectors $(\boldsymbol{q}_a, \boldsymbol{q}_b, \boldsymbol{q}_c)$. Each eigenvector defines a principal direction of elongation of the cell doublet, to which we can associate three principal radii $(a, b, c) = (\sqrt{w_a}, \sqrt{w_b}, \sqrt{w_c})$. We consider their relative values $(a/b, 1, c/b)$ to quantify the aspect ratio of the doublet, in a way that is invariant with respect to the doublet's total size. In Figs. 2c and 4g, we correlate the directions of elongations $\boldsymbol{q}_i$ with the rotation vector $\boldsymbol{\omega}$ by computing the following quantities:

$$C_{\boldsymbol{\omega},\boldsymbol{q}_i} = \left\langle \frac{3}{2}\left(\frac{\boldsymbol{\omega}}{||\boldsymbol{\omega}||} \cdot \boldsymbol{q}_i\right)^2 - \frac{1}{2} \right\rangle \ . \tag{7}$$

This quantity is invariant by $\boldsymbol{q}_i \to -\boldsymbol{q}_i$, reflecting the nematic property of an elongation axis. It is equal to 1 if the vectors are always parallel, $-1/2$ if they are always perpendicular, and 0 if they have uniformly distributed random orientations.

In Fig. 2c, we also correlate the longest elongation vector $\boldsymbol{q}_a$ with the vector $\boldsymbol{r}_{12}$ joining the center of masses of the cells, using the following formula:

$$C_{\boldsymbol{r}_{12},\boldsymbol{q}_a} = \left\langle \left| \frac{\boldsymbol{r}_{12}}{||\boldsymbol{r}_{12}||} \cdot \boldsymbol{q}_a \right| \right\rangle \ . \tag{8}$$

### 4 Interface shape analysis

In this section we explain how the interface between the cells is extracted from segmented meshes, how the interface shape is analysed quantitatively and how it is correlated with the doublet rotation.

#### 4.1 Interface extraction

For every frame $t$ of every segmented movie, the segmentation process produces meshes with vertices $\{\boldsymbol{v}_{1i} = \{v_{1ix}, v_{1iy}, v_{1iz}\}\}, \{\boldsymbol{v}_{2i}\}$ representing the membranes of cell 1 and 2 of the cell doublet. As discussed above, we define a distance threshold of $d_{th} = 10$ pixels and select all vertices (from both cells) that are less than a distance $d_{th}$ away from the other cell, in order to obtain the vertices of the interface $\{\boldsymbol{v}_{\text{int},i}$ for $i \in \{1, ..., N\}\}$. We then use the method of principal component analysis (PCA) to find the direction which minimizes the variance of the coordinates of the points. We start by computing the center of mass of the interface vertices $\boldsymbol{C} = \frac{1}{N}\sum_i \boldsymbol{v}_{\text{int},i}$. In order to reduce the dependency of $\boldsymbol{C}$ on the discreteness of the meshes, we define another center $\boldsymbol{C}'$ which is the projection of $\boldsymbol{C}$ onto the $\boldsymbol{r}_{12}$ axis:

$$\boldsymbol{C}' = \boldsymbol{r}_1 + \boldsymbol{r}_{12}\frac{\boldsymbol{r}_{12}\cdot(\boldsymbol{C}-\boldsymbol{r}_1)}{||\boldsymbol{r}_{12}||^2} \ , \tag{9}$$

where $\boldsymbol{r}_1$ is the center of mass of cell 1. We then do the PCA by performing a singular value decomposition of the matrix $M_{ji} = v_{\text{int},ij} - C'_j$ with $j \in \{x, y, z\}$ and $i = \{1, ..., N\}$ [2]. This generates three orthonormal vectors that are aligned according to the principal directions of elongations of the point set $\{\boldsymbol{v}_{\text{int},i}\}$. We define the plane of the interface as the plane normal to the unit vector $\boldsymbol{N}_{12}$ which is aligned to the direction of minimum elongation, and going from cell 1 towards cell 2. The other two vectors $(\boldsymbol{e}_X, \boldsymbol{e}_Y)$ define a two-dimensional basis in the interface plane. This allows us to define a set of local coordinates $(X, Y, H)$ such that the vertices of the interface have positions:

$$\boldsymbol{v}_{\text{int},i} = \boldsymbol{C}' + X_i\boldsymbol{e}_X + Y_i\boldsymbol{e}_Y + H_i\boldsymbol{N}_{12} \ . \tag{10}$$

The first step to analyse the shape of the interface is to analyse its shape in the $(\boldsymbol{e}_X, \boldsymbol{e}_Y)$ plane. For this we construct a tensor of rank 2 $\mathbf{Q}^{2d}$ defined as:

$$Q^{2d}_{\alpha\beta} = \frac{1}{N}\sum_{i=1}^{N}(v_{\text{int},i,\alpha} - C'_\alpha)(v_{\text{int},i,\beta} - C'_\beta) \text{ with } \alpha, \beta \text{ in } \{X, Y\} \ . \tag{11}$$

This symmetric tensor can be diagonalized, which gives two eigenvalues $(w_1, w_2)$ $(w_1 \geq w_2)$ associated to two unit eigenvectors $(\boldsymbol{\kappa}_1, \boldsymbol{\kappa}_2)$.

In the case of an ellipse of large radius $\lambda R_e$ oriented along the axis $x$ and small radius $R_e/\lambda$ oriented along the axis $y$, in the continuous limit for points homogeneously distributed, we have (here for the $Q^{2d}_{xx}$ component):

$$Q^{2d(\text{ellipse})}_{xx} = \frac{1}{\pi R_e^2}\int_{x=-\lambda R_e}^{\lambda R_e} dx \int_{y=-\frac{R_e}{\lambda}\sqrt{1-\frac{x^2}{\lambda^2 R_e^2}}}^{\frac{R_e}{\lambda}\sqrt{1-\frac{x^2}{\lambda^2 R_e^2}}} dy x^2 = \frac{\lambda^2 R_e^2}{\pi}\int_{x=-1}^{1} dx \int_{y=-\sqrt{1-x^2}}^{\sqrt{1-x^2}} dy x^2 = \frac{\lambda^2 R_e^2}{4} \ . \tag{12}$$

The other component can be computed similarly, which leads to:

$$\boldsymbol{Q}^{2d(\text{ellipse})} = \begin{pmatrix} \frac{\lambda^2 R_e^2}{4} & 0 \\ 0 & \frac{R_e^2}{4\lambda^2} \end{pmatrix} \ , \tag{13}$$

from which one can show that the eigenvalues $w_1$, $w_2$ are related to $\lambda$ and $R_e$ by:

$$R_e = 2\left(w_1 w_2\right)^{1/4}, \ \lambda = \left(\frac{w_1}{w_2}\right)^{1/4} \ . \tag{14}$$

We use these formulas to compute an effective radius $R_e$ and an effective aspect ratio $\lambda \geq 1$ for the interface. Then, we compute the planar coordinates $(U_{p,i}, V_{p,i})$ of the interface points in the reference frame defined by the perpendicular eigenvectors $(\boldsymbol{\kappa}_1, \boldsymbol{\kappa}_2)$:

$$\begin{aligned} U_{p,i} &= X_i \kappa_{1x} + Y_i \kappa_{1y} \ , \\ V_{p,i} &= X_i \kappa_{2x} + Y_i \kappa_{2y} \ . \end{aligned} \tag{15}$$

We then bring back the interface onto a dimensionless disk of unit radius by rescaling the coordinates $(U_{p,i}, V_{p,i})$ into $(U'_{p,i}, V'_{p,i})$ given by:

$$\begin{aligned} U'_{p,i} &= \frac{U_{p,i}}{R_e \lambda} \ , \\ V'_{p,i} &= \frac{\lambda V_{p,i}}{R_e} \ . \end{aligned} \tag{16}$$

Finally, we express the rescaled coordinates in the original reference frame as $(\tilde{X}_i, \tilde{Y}_i, \tilde{H}_i)$:

$$\begin{aligned} \tilde{X}_i &= U'_{p,i} \kappa_{1x} + V'_{p,i} \kappa_{2x} \ , \\ \tilde{Y}_i &= U'_{p,i} \kappa_{1y} + V'_{p,i} \kappa_{2y} \ , \\ \tilde{H}_i &= \frac{H_i}{R_e} \ . \end{aligned} \tag{17}$$

Note that we also divided $H_i$ by $R_e$ to preserve the aspect ratio of the interface in the vertical dimension. The mode decomposition of the interface is then performed on the reduced coordinates $(\tilde{X}_i, \tilde{Y}_i, \tilde{H}_i)$.

#### 4.2 Mode decomposition

On a unit disk $D_1$ defined by $(x, y) \in \mathbb{R}^2$ such that $x^2 + y^2 \leq 1$, we want to find a function $H(x, y)$ that is smooth and passes as close as possible to the experimental points $(\tilde{X}_i, \tilde{Y}_i, \tilde{H}_i)$. We choose to pick $H$ among the set of polynomials of degree $n \leq 3$, which represents a good compromise between the quality of the fit to the experimental points and the complexity of the description. We therefore define the interface polynomial $P$ which satisfies the following linear least square problem:

$$P(x, y) = \sum_{i=0}^{3} \sum_{j=0}^{i} a_{ij} x^{i-j} y^j \text{ with } \{a_{ij}\} \text{ such that } \sum_{k=1}^{N} \left( P(\tilde{X}_k, \tilde{Y}_k) - \tilde{H}_k \right)^2 \text{ is minimized .} \tag{18}$$

Once the polynomial $P$ is found, we seek to represent it in a basis that is well suited for the study of the symmetries of the cell-cell interface shape. We propose the following basis:

$$
\begin{aligned}
&\text{Mode of degree 0: } \frac{1}{\sqrt{\pi}} \ . \\
&\text{Modes of degree 1: } \frac{2x}{\sqrt{\pi}}, \ \frac{2y}{\sqrt{\pi}} \ . \\
&\text{Modes of degree 2: } B = 2\sqrt{\frac{3}{\pi}}\left(x^2+y^2-\frac{1}{2}\right) \ , \\
&\qquad S_1 = \sqrt{\frac{6}{\pi}}(x^2-y^2), \ S_2 = 2\sqrt{\frac{6}{\pi}}xy \ . \\
&\text{Modes of degree 3: } T_{f1} = 2\sqrt{\frac{2}{\pi}}x(x^2-3y^2), \ T_{f2} = 2\sqrt{\frac{2}{\pi}}y(y^2-3x^2) \ , \\
&\qquad Y_{y1} = 6\sqrt{\frac{2}{\pi}}x\left(x^2+y^2-\frac{2}{3}\right), \ Y_{y2} = 6\sqrt{\frac{2}{\pi}}y\left(y^2+x^2-\frac{2}{3}\right) \ .
\end{aligned}
\tag{19}
$$

which, up to a normalisation, corresponds to Zernike polynomials of order up to 3 [3]. This basis is orthonormal on the unit disk with the following definition of the scalar product:

$$
A \cdot B = \int_{D_1} dxdy A(x,y)B(x,y) \tag{20}
$$

Also, special care has been taken to construct basis vectors with specific symmetries. The $S_1$ and $S_2$ polynomials are saddle-nodes and belong to the $D_{2d}$ symmetry group. $B$ is an isotropic mode with a “bowl” shape belonging to the $C_{\infty v}$ group. $T_{f1}$ and $T_{f2}$ have three-fold symmetry and belong to the $D_{3d}$ group. Finally, $Y_{y1}$ and $Y_{y2}$, the “yin-yang” modes, belong to the $C_{2h}$ group (Supplementary Table 1).

The interface polynomial $P$ can be projected in the basis and reads:

$$
\begin{aligned}
P(x,y) = \frac{a_0}{\sqrt{\pi}} + a_1\frac{2x}{\sqrt{\pi}} + a_2\frac{2y}{\sqrt{\pi}} + a_3 S_1(x,y) + a_4 S_2(x,y) + a_5 B(x,y) \\
+ a_6 T_{f1}(x,y) + a_7 T_{f2}(x,y) + a_8 Y_{y1}(x,y) + a_9 Y_{y2}(x,y) \ .
\end{aligned}
\tag{21}
$$

The coefficients $a_i$ contains all the information about the shape of the interface (assuming that it is well represented by the polynomial $P$). For instance one can compute the average squared deflection of the interface $\langle \tilde{H}^2 \rangle$:

$$
\langle \tilde{H}^2 \rangle = \frac{1}{\pi}\int_{D_1} dxdy P(x,y)^2 = \frac{1}{\pi}\sum_{i=0}^{9} a_i^2 \ , \tag{22}
$$

The average deflection of the interface in real units can be expressed as:

$$
\sqrt{\langle H^2 \rangle} = \frac{R_e}{\sqrt{\pi}}\left(\sum_{i=0}^{9} a_i^2\right)^{1/2} \ , \tag{23}
$$

with $R_e$ the radius of the interface computed in Eq. 14. Each factor $a_i^2$ can be seen as a quantification of the weight of the mode $i$ in the interface deflection. Particularly, we are interested in the

| Mode | Symmetry group | Invariant symmetry operation |
| --- | --- | --- |
| Bowl 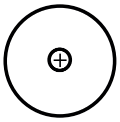 | $C_{\infty v}$ | $c_{\infty}^{Z}$, $\sigma_{\infty}^{X,Y}$ |
| Yin-yang 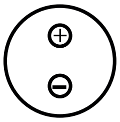 | $C_{2h}$ | $c_2^X$, $\sigma^X$, $I$ |
| Three-fold 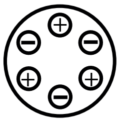 | $D_{3d}$ | $I$, $\sigma^X$, $\sigma^a$, $\sigma^b$, $c_2^X$ ,$c_2^a$, $c_2^b$, $c_3^Z$, $c_{-3}^Z$, $s_6^Z$, $s_{-6}^Z$ |
| Saddle-node 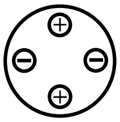 | $D_{2d}$ | $\sigma^X$, $\sigma^Y$, $c_2^c$, $c_2^d$, $c_2^Z$, $s_4^Z$, $s_{-4}^Z$ |

**Supplementary Table 1:** Classification of interface deformation modes and associated symmetries. See Ext. Fig. 4b for a schematic of axis orientations. The $Z$ axis corresponds to the axis along the vector $\mathbf{N}_{12}$. $I$ indicates the inversion symmetry. $c_2^\alpha$, $c_3^\alpha$ $c_{-3}^\alpha$ are the rotations by $\pi$, $2\pi/3$ and $-2\pi/3$ around the axis given by $\alpha$. $c_\infty^Z$ is the set of rotations by any angle around the $Z$ axis. $\sigma^\alpha$ is the mirror symmetry with respect to the plane normal to the $\alpha$ direction, $\sigma_\infty^{X,Y}$ denotes the set of mirror symmetries with respect to a plane containing the $Z$ axis. $s_4^Z$, $s_{-4}^Z$, $s_6^Z$ and $s_{-6}^Z$ are improper rotations of $\pi/2$, $-\pi/2$, $\pi/3$ and $-\pi/3$ around the Z axis. The vectors $\mathbf{a}$, $\mathbf{b}$, $\mathbf{c}$, $\mathbf{d}$ are obtained by rotation of angle $-2\pi/3$, $2\pi/3$, $\pi/4$, $-\pi/4$ of the vector $\mathbf{e}_X$ along the vector $\mathbf{e}_Z$ (Ext. Fig. 4b).

weights of the "saddle-node", "bowl", "three-fold" and "yin-yang" modes:

$$\begin{aligned} W_S &= a_3^2 + a_4^2 \\ W_B &= a_5^2 \\ W_T &= a_6^2 + a_7^2 \\ W_Y &= a_8^2 + a_9^2 \end{aligned} \qquad (24)$$

We ignore the weights of the remaining modes, of order zero and one, which have vanishing or small amplitude. Indeed these modes describe a deviation of the interface towards a new plane, which we expect is removed to a large extent by the plane fitting procedure described above. We then divide each term in Eq. 24 by the sum $W_S + W_B + W_T + W_Y$ to generate relative mode amplitudes that are shown in Fig. 2g, Fig. 4f,j and Ext. Fig. 6d.

#### 4.3 Mode directions and correlation with rotation

In order to correlate the modes of the interface with the rotation of the doublet, we need first to identify their principal directions in the plane $(X, Y)$ as a function of the coefficients $a_i$. These directions can only be defined up to the rotational symmetry of each mode, as illustrated on Ext. Fig. 4a, and also exist only relatively to the oriented plane of the interface $\boldsymbol{N}_{12}$, going from cell 1 to cell 2. We define the normalized vectors $\boldsymbol{Y}_{12}$, $\boldsymbol{S}_{12}$ and $\boldsymbol{T}_{12}$ associated respectively to the yin-yang, saddle-node and three-fold modes as follows:

$$\begin{aligned}
\boldsymbol{Y}_{12} &= \frac{a_8\boldsymbol{e}_X + a_9\boldsymbol{e}_Y}{\sqrt{a_8^2+a_9^2}} \ , \\
\boldsymbol{S}_{12} &= \cos(\alpha)\boldsymbol{e}_X + \sin(\alpha)\boldsymbol{e}_Y \text{ with } \alpha = \frac{1}{2}\arctan\left(a_3, a_4\right) + n\pi \ , \\
\boldsymbol{T}_{12} &= \cos(\alpha)\boldsymbol{e}_X + \sin(\alpha)\boldsymbol{e}_Y \text{ with } \alpha = \frac{1}{3}\arctan\left(-a_6, a_7\right) + \frac{2n\pi}{3} \ ,
\end{aligned} \tag{25}$$

where $\arctan(x, y)$ is the arc tangent of $y/x$ (taking into account in which quadrant the point is located), and $n$ is an arbitrary integer, representative of the rotational symmetries of the saddle-node and three-fold modes. The vectors are transformed in the following way under a switch of cell indices:

$$\begin{aligned}
\boldsymbol{Y}_{21} &= -\boldsymbol{Y}_{12} \text{ (sign change)}, \\
\boldsymbol{S}_{21} &= \boldsymbol{N}_{12} \times \boldsymbol{S}_{12} \text{ } (90^\circ \text{ rotation}), \\
\boldsymbol{T}_{21} &= -\boldsymbol{T}_{12} \text{ (sign change)}.
\end{aligned} \tag{26}$$

Now, the correlation between these vectors and the rotation vector $\boldsymbol{\omega}$ of the doublet must be constructed in such a way that it is independent of the arbitrary choice of the indices of the cells, and is also invariant under change of the variable $n$ describing the rotation symmetries of individual modes. To this effect, we introduce the vector $\boldsymbol{u}_{12} = \boldsymbol{N}_{12} \times \boldsymbol{\omega}/||\boldsymbol{N}_{12} \times \boldsymbol{\omega}||$ and define the correlations as follows:

$$\begin{aligned}
C_{\boldsymbol{\omega},\boldsymbol{Y}} &= \langle \boldsymbol{Y}_{12} \cdot \boldsymbol{u}_{12} \rangle \ , \\
C_{\boldsymbol{\omega},\boldsymbol{S}} &= \langle \cos\left(4 \arccos\left(\boldsymbol{S}_{12} \cdot \boldsymbol{u}_{12}\right)\right) \rangle \ , \\
C_{\boldsymbol{\omega},\boldsymbol{T}} &= \langle \cos\left(3 \arccos\left(\boldsymbol{T}_{12} \cdot \boldsymbol{u}_{12}\right)\right) \rangle \ .
\end{aligned} \tag{27}$$

These three formulas are used to compute the correlations presented in Figs. 2h and 4g. Note the necessity of introducing a 4-th degree correlation function for $C_{\boldsymbol{\omega},\boldsymbol{S}}$ because of the peculiar way it transforms itself under a cell index switch. Indeed we have:

$$\arccos(\boldsymbol{S}_{21} \cdot \boldsymbol{u}_{21}) = \arccos(-(\boldsymbol{N}_{12} \times \boldsymbol{S}_{12}) \cdot \boldsymbol{u}_{12}) \ . \tag{28}$$

In the reference frame $(\boldsymbol{u}_{12}, \boldsymbol{N}_{12} \times \boldsymbol{u}_{12}, \boldsymbol{N}_{12})$, the coordinates of $\boldsymbol{S}_{12}$ are $(\cos(\theta_S), \sin(\theta_S), 0)$. $\theta_S$ can be defined to be in the interval $[-\pi, \pi]$, in which case we have:

$$\begin{aligned}
\arccos(\boldsymbol{S}_{12} \cdot \boldsymbol{u}_{12}) &= \arccos(\cos(\theta_S)) = \pm\theta_S, \ (+ \text{ if } \theta_S \in [0, \pi]) \ , \\
\arccos(\boldsymbol{S}_{21} \cdot \boldsymbol{u}_{21}) &= \arccos(\sin(\theta_S)) = \begin{cases} (\theta_S - \frac{\pi}{2}, & \text{if } \theta_S \in \left[\frac{\pi}{2}, \pi\right], \\ -\left(\theta_S - \frac{\pi}{2}\right), & \text{if } \theta_S \in \left[-\frac{\pi}{2}, \frac{\pi}{2}\right] \\ \theta_S + \frac{3\pi}{2}, & \text{if } \theta_S \in \left[-\pi, -\frac{\pi}{2}\right] \end{cases} \ ,
\end{aligned} \tag{29}$$

where the second line is obtained using Eq. 28. From Eq. 29, we see that in order to get an invariant quantity after index switch, one need to multiply by (at least) 4 and take the cosine:

$$\cos(4 \arccos(\boldsymbol{S}_{21} \cdot \boldsymbol{u}_{21})) = \cos(4 \arccos(\boldsymbol{S}_{12} \cdot \boldsymbol{u}_{12})) \ \forall \theta_S \in [-\pi, \pi] \ . \tag{30}$$

Concerning the bowl mode, we rename its coefficient $a_5$ into $B_{12}$, a signed scalar quantity which indicates the direction of it with respect to the vector $\boldsymbol{N}_{12}$ (see Ext. Fig. 4a).

##### 4.4 Doublet and interface symmetries

In order to apply the Curie principle, we need to determine symmetry point groups of the doublet which are compatible with the interface deformation. In Table 2, we list a set of possible symmetry groups to which the cortical myosin distribution of the doublet can belong, and by comparison of the invariant transformations also leaving interface modes invariant (Table 1), we find compatible interface deformations (Fig. 4j).

| Example | Symmetry group | Invariant symmetry operations | Compatible interface deformation mode |
|---|---|---|---|
| 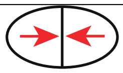 | $D_{\infty h}$ | $c_\infty^Z$, $c_2^{X,Y}$, $s_\infty^Z$, $\sigma^Z$, $\sigma_\infty^{X,Y}$, $I$ | None |
| 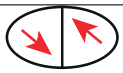 | $C_{2h}$ | $c_2^X$, $\sigma^X$, $I$ | $C_{2h}$, $D_{3d}$ |
| 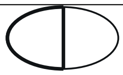 | $C_{\infty v}$ | $c_\infty^Z$, $\sigma_\infty^{X,Y}$ | $C_{\infty v}$ |
| 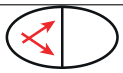 | $C_{2v}$ | $c_2^Z$, $\sigma^X$, $\sigma^Y$ | $C_{\infty v}$, $D_{2d}$ |
| 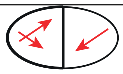 | $C_s$ | $\sigma^X$ | $C_{\infty v}$, $D_{3d}$, $C_{2h}$, $D_{2d}$ |

**Supplementary Table 2:** Classification of doublet symmetry properties and associated interface deformation modes. $I$ indicates the inversion symmetry. $c_2^\alpha$ the rotation by $\pi$ around the axis given by $\alpha$. $\sigma^\alpha$ the mirror symmetry with respect to the plane normal to the $\alpha$ direction. $c_\infty^Z$ denotes the set of rotation symmetry by any angle around the $Z$ axis. $\sigma_\infty^{X,Y}$ denotes the set of mirror symmetries with respect to any plane containing the $Z$ axis. $s_\infty^Z$ denotes the set of improper rotations by any angle around the $Z$ axis. $c_2^{X,Y}$ denotes the set of rotation by $\pi$ around any axis contained in the $X, Y$ plane. See Ext. Fig. 4b for a schematic of space orientation. Compatible deformation modes are obtained using the Curie principle and Table 1.

#### 5 Analysis of cortical myosin fluorescence intensity profiles

In this section, we describe how a polarity vector and nematic tensor are extracted for each cell from the distribution of cortical myosin, as measured from the MRLC fluorescence intensity profiles.

##### 5.1 Obtention of cortical myosin profiles

The first step of the polarity analysis process is to extract the myosin fluorescence signal from the microscopy images and associate it to the vertices $\{\boldsymbol{v}_{1i}\}, \{\boldsymbol{v}_{2i}\}$ of the segmented cell membranes. One challenge to overcome is that typical segmentation errors often put the mesh slightly away from the real cell membrane location, so that the vertices are not perfectly colocated with pixels belonging to the cell cortex. To solve for this issue, we sample voxels in a rectangular prism around each vertex as described on Ext. Fig. 4c. The larger the sampling rectangular prism, the higher the chance there is to correct for a mismatch of the vertex position relative to the cell cortex, but a larger sampling region also captures dark voxels outside of the cell, which artificially decreases the overall signal. To avoid this issue, we choose to average the values of the $n$ brightest pixels in the sampling rectangular prism. In this study we used a rectangular window of 11x11x3 voxels in size and a fraction of $n = 20\%$ of the brightest voxels within the window.

In order to compensate for the background signal due to the cytosolic concentration of myosin, which we assume does not play a role in the modulation of the cortical tension, we average the signal of all the voxels inside of the mesh and subtracts this value to the signal on the vertices. Furthermore, since the signal observed at the vertices of the cell-cell interface most probably is a superposition of the signal coming from both cells, we divide by two the value of the measured signal, before subtracting the cytosolic value.

#### 5.2 Correction of signal intensity profiles away from the microscope

The $z$ axis of the coordinate system of the fluorescence images is the axis that measures the distance to the microscope. The larger the $z$ values of a voxel, the more materials there is between this voxel and the microscope. This can lead to an artificial bias in the signal measured. In order to quantify this, we display the membrane myosin signal (after attaching it to vertices following the procedure in section 5.1) of individual doublets along $x$, $y$, and $z$ (Ext. Fig. 4d). We typically observe a decrease along $z$, but not along $x$ and $y$. We measure this bias more precisely by fitting exponential profiles to the time averaged signals of each doublets:

$$c^{\text{fit}}(x) = A_x \exp\left(\lambda_x x\right) \ , \ c^{\text{fit}}(y) = A_y \exp\left(\lambda_y y\right) \ , \ c^{\text{fit}}(z) = A_z \exp\left(\lambda_z z\right) \ . \tag{31}$$

We extract $\lambda_x^i$, $\lambda_y^i$, $\lambda_z^i$ for each doublet $i$, and we see that only the $\lambda_z$ coefficients are significantly negative (Ext. Fig. 4e). We therefore constructed a corrected myosin signal $c^i(\{x, y, z\}, t)$ for each doublet $i$:

$$c^i(\{x, y, z\}, t) = \exp\left(-\lambda_z^i z\right) c^{i,\text{measured}}(\{x, y, z\}, t) \ . \tag{32}$$

We perform this correction on all the voxels of all the microscopy images. We then re-compute the myosin signal on vertices (section 5.1) and continue the analysis. The correction made here is valid under the hypothesis that the signal decrease observed along $z$ is due to optical effects and has no biological origin.

#### 5.3 Computation of polarity from cortical myosin distribution

We wish to quantify the heterogeneity of the cortical myosin distribution. We first define the cell surface center of mass

$$\boldsymbol{r}_g = \frac{1}{A} \int dS \mathbf{r} \ , \tag{33}$$

with $A$ the surface area. We then construct a polarity vector, originating from $\boldsymbol{r}_g$, that points towards the region of higher myosin signal. There can be multiple ways to define such a polarity vector. On a surface $S$, with a myosin signal $c(\boldsymbol{r})$, we adopt the following definition of $\boldsymbol{p}$:

$$\boldsymbol{p} = \frac{\int_S dS' \frac{\Delta \boldsymbol{r}}{||\Delta \boldsymbol{r}||} c(\boldsymbol{r})}{\int_S dS' c(\boldsymbol{r})} \ , \tag{34}$$

with $\Delta \boldsymbol{r} = \boldsymbol{r} - \boldsymbol{r}_g$, and $dS'$ is the projection on a unit sphere centered around $\boldsymbol{r}_g$, of the area element $dS$ around a point $\boldsymbol{r}$ on the surface. Defined in this way, $\boldsymbol{p}$ is invariant by a rescaling of the signal $c(\boldsymbol{r})$, and a rescaling of space. Also, if all the signal is concentrated in an infinitely small spot $\boldsymbol{r}_c$ ($c(\boldsymbol{r}) = \alpha\delta(\boldsymbol{r} - \boldsymbol{r}_c)$) one can show that we have $||\boldsymbol{p}|| = 1$. Integrating on a unit sphere, instead of on the actual surface $S$, reduces the dependency of the polarity on cell elongation. On a triangle mesh, assuming a homogeneous signal $c(\boldsymbol{r}) = c_i$ on each triangle $t_i$ with area $A_i$, we compute first the center of mass $\boldsymbol{r}_g$ as:

$$\boldsymbol{r}_g = \frac{\sum_{i=1}^{N_t} A_i \boldsymbol{r}_i}{A} \text{ with } \boldsymbol{r}_i = \frac{1}{A_i}\int_{t_i} dS \text{ and } A = \sum_i A_i \boldsymbol{r} \ . \tag{35}$$

We then compute the coordinates $(\theta_i, \phi_i)$ of the center of each triangle, in the spherical coordinate system defined as:

$$\begin{aligned} \frac{\Delta r_x}{||\Delta\boldsymbol{r}||} &= \sin(\theta)\cos(\phi) \ , \\ \frac{\Delta r_y}{||\Delta\boldsymbol{r}||} &= \sin(\theta)\sin(\phi) \ , \\ \frac{\Delta r_z}{||\Delta\boldsymbol{r}||} &= \cos(\theta) \ . \end{aligned} \tag{36}$$

We then create equally spaced bins in $\theta$, $\phi$ and we compute the signal in each bin, which is the average of the signal $c_i$ of all the triangles $t_i$ whose projected center has its coordinates $(\theta_i, \phi_i)$ inside the bin. If a bin contains no triangle, we take the signal of the closest projected triangle center. We finally compute the polarity using the following integral, discretised on the bins:

$$\begin{aligned} p_x &= \frac{1}{\langle c\rangle}\int_{\theta,\phi} d\theta d\phi \sin(\theta)\sin(\theta)\cos(\phi)c(\theta,\phi) \ , \\ p_y &= \frac{1}{\langle c\rangle}\int_{\theta,\phi} d\theta d\phi \sin(\theta)\sin(\theta)\sin(\phi)c(\theta,\phi) \ , \\ p_z &= \frac{1}{\langle c\rangle}\int_{\theta,\phi} d\theta d\phi \sin(\theta)\cos(\theta)c(\theta,\phi) \ , \\ \langle c\rangle &= \int_{\theta,\phi} d\theta d\phi \sin(\theta)c(\theta,\phi) \ . \end{aligned} \tag{37}$$

#### 5.4 Polarity orientiations in a cell doublet

In a rotating doublet, we describe the orientations of the polarities $\boldsymbol{p}_1$ and $\boldsymbol{p}_2$ of the two cells by introducing a set of angles $\alpha_1$, $\alpha_2$ and $\beta$. Using the vector $\boldsymbol{r}_{12}$ joining the center of mass of cell 1 to the center of mass of cell 2, we define $\alpha_i$ as:

$$\begin{aligned} \alpha_1 &= \arccos\left(\frac{\boldsymbol{p}_1}{||\boldsymbol{p}_1||}\cdot\frac{\boldsymbol{r}_{12}}{||\boldsymbol{r}_{12}||}\right) \ , \\ \alpha_2 &= \arccos\left(\frac{\boldsymbol{p}_2}{||\boldsymbol{p}_2||}\cdot\frac{-\boldsymbol{r}_{12}}{||\boldsymbol{r}_{12}||}\right) \ . \end{aligned} \tag{38}$$

These are the angles between the polarities and the axis $\boldsymbol{r}_{12}$ of the doublet (Fig. 3f,g). In order to quantify how the polarities are organised in the plane perpendicular to $\boldsymbol{r}_{12}$, we first introduce projected polarities $\boldsymbol{p}_i^{\perp}$:

$$\boldsymbol{p}_i^{\perp} = \boldsymbol{p}_i - (\boldsymbol{p}_i\cdot\boldsymbol{r}_{12})\frac{\boldsymbol{r}_{12}}{||\boldsymbol{r}_{12}||^2} \ . \tag{39}$$

We then define a 2D basis $(\boldsymbol{u}_1, \boldsymbol{u}_2)$ around $\boldsymbol{p}_1^\perp$:

$$\boldsymbol{u}_1 = \frac{\boldsymbol{p}_1^\perp}{||\boldsymbol{p}_1^\perp||} \ , \ \boldsymbol{u}_2 = \frac{\boldsymbol{r}_{12}}{||\boldsymbol{r}_{12}||} \times \boldsymbol{u}_1 \ . \tag{40}$$

We define $\beta$ as the angle of $\boldsymbol{p}_2^\perp$ in this basis:

$$\cos(\beta) = \frac{\boldsymbol{p}_2^\perp}{||\boldsymbol{p}_2^\perp||} \cdot \boldsymbol{u}_1 \ , \ \sin(\beta) = \frac{\boldsymbol{p}_2^\perp}{||\boldsymbol{p}_2^\perp||} \cdot \boldsymbol{u}_2 \ . \tag{41}$$

When the vectors $\boldsymbol{p}_1$, $\boldsymbol{p}_2$ and $\boldsymbol{r}_{12}$ are coplanar, $\beta$ is equal to 0 or $\pi$. $\beta = 0$ means that $\boldsymbol{p}_2^\perp$ and $\boldsymbol{p}_1^\perp$ are parallel in the same direction, and $\beta = \pi$ means that $\boldsymbol{p}_2^\perp$ and $\boldsymbol{p}_1^\perp$ are parallel in the opposite direction. The distribution of $\beta$ is shown on Fig. 3h for all time points of all doublets. We define the average of the distribution $\{\beta_i\}$ for $N$ doublets in a way that is suited to a distribution of angles:

$$\langle\beta\rangle = \arctan\left(\frac{1}{N}\sum_i \cos(\beta_i), \frac{1}{N}\sum_i \sin(\beta_i))\right) \ , \tag{42}$$

where the arctan function is defined in the same way as in Eq.25. The corresponding average is shown as a red line in Fig. 3h, and is close to $\pi$.

#### 5.5 Correlation of myosin polarity with rotation vector and interface modes

In order to test the assumption that the doublet rotation $\boldsymbol{\omega}$ is correlated with the orientation of the cell polarities $\boldsymbol{p}_1$ and $\boldsymbol{p}_2$, we need to construct a parameter that quantifies this correlation. We choose to correlate $\boldsymbol{\omega}$ with $\boldsymbol{r}_{12} \times (\boldsymbol{p}_1 - \boldsymbol{p}_2)$, as these two quantities are pseudovectors and are invariant under an exchange of cell indices $1 \leftrightarrow 2$ (Fig. 3i):

$$C(\boldsymbol{\omega}, \boldsymbol{r}_{12} \times (\boldsymbol{p}_1 - \boldsymbol{p}_2)) \text{ with } C(\boldsymbol{a}, \boldsymbol{b}) = \left\langle \frac{\boldsymbol{a}}{||\boldsymbol{a}||} \cdot \frac{\boldsymbol{b}}{||\boldsymbol{b}||} \right\rangle \ . \tag{43}$$

This correlation takes values between $-1$ and 1 and is invariant under a change of cell indices. Additionally, we can correlate the polarity orientations with the "yin-yang" deformation mode of the interface defined in Eq. 25 (Fig. 3j):

$$C(\boldsymbol{Y}_{12}, \boldsymbol{p}_1 - \boldsymbol{p}_2) \ , \tag{44}$$

with the same definition of $C(\boldsymbol{a}, \boldsymbol{b})$ as in Eq. 43. Again this quantity is invariant under a change of cell indices.

#### 5.6 Generation of time averaged polarity maps

In Figs. 3, 4 of the main text, we generate maps in spherical coordinates of the averaged cortical myosin distribution on the cell surface. We average the signal on all time points and on all the cells. We define two orthogonal reference frames that are locally defined around each cell $i$ and can be used to represent the signal. The first kind of reference frame is constructed around $\boldsymbol{\omega}_i$ (the rotation vector of the doublet containing cell $i$) and $\boldsymbol{r}_{in_i}$ ($n_i$ being the other cell in the doublet containing cell $i$), and the second kind is constructed around the polarity vector itself $\boldsymbol{p}_i$ and $\boldsymbol{r}_{in_i}$:

$$\begin{aligned}(\boldsymbol{e}_x^{(i)}, \boldsymbol{e}_y^{(i)}, \boldsymbol{e}_z^{(i)})_1 &= \left(\frac{\boldsymbol{\omega}_i}{||\boldsymbol{\omega}_i||}, \boldsymbol{e}_z^{(i)} \times \boldsymbol{e}_x^{(i)}, \frac{\boldsymbol{r}_{in_i}}{||\boldsymbol{r}_{in_i}||}\right) \ , \\ (\boldsymbol{e}_x^{(i)}, \boldsymbol{e}_y^{(i)}, \boldsymbol{e}_z^{(i)})_2 &= \left(\frac{\boldsymbol{p}_i}{||\boldsymbol{p}_i||} \times \frac{\boldsymbol{r}_{in_i}}{||\boldsymbol{r}_{in_i}||}, \boldsymbol{e}_z^{(i)} \times \boldsymbol{e}_x^{(i)}, \frac{\boldsymbol{p}_i}{||\boldsymbol{p}_i||}\right) \ .\end{aligned} \tag{45}$$

We use spherical coordinates to define a couple of angles $(\theta_j, \phi_j)$ for each vertex $j$ of a cell $i$:

$$\frac{\boldsymbol{r}_j - \boldsymbol{r}_g^{(i)}}{||\boldsymbol{r}_j - \boldsymbol{r}_g^{(i)}||} = \cos(\phi_j)\sin(\theta_j)\boldsymbol{e}_x^{(i)} + \sin(\phi_j)\sin(\theta_j)\boldsymbol{e}_y^{(i)} + \cos(\theta_j)\boldsymbol{e}_z^{(i)} \ , \tag{46}$$

with $\boldsymbol{r}_g^{(i)}$ the center of mass of cell $i$. We make equally spaced bins in $\theta$, $\phi$ and we accumulate the normalized myosin signal of each triangle of each cell (for every doublet and every time point) into the bins to finally construct an average myosin signal profile, which is represented on Figs. 3 and 4 of the main text. We normalize the myosin signal of a given time point of a given cell by rescaling it such that its average value on the unit sphere ($\theta \in [0, \pi]$, $\phi \in [-\pi, \pi]$) is 1. The average value on the unit sphere is obtained from discretisation of the following averaging operation for the field $I$:

$$\langle I \rangle = \frac{1}{4\pi}\int_0^{2\pi} d\phi \int_0^{\pi} d\theta \sin\theta I \ . \tag{47}$$

#### 5.7 Second order description of the myosin signal heterogeneity

The polarity vector $\boldsymbol{p}$ can be considered a lowest order characterisation of the heterogeneity of the myosin profile around the cell. To go further, one can calculate the nematic tensor associated to the cortical myosin fluorescence intensity profile $s(\boldsymbol{r})$:

$$Q_{\alpha\beta} = \frac{\int_S dS(\overline{r}_\alpha \overline{r}_\beta - \frac{1}{3}\delta_{\alpha\beta}(\overline{r}_\gamma \overline{r}_\gamma))s(\boldsymbol{r})}{\int_S dS s(\boldsymbol{r})} \text{ with } \overline{\boldsymbol{r}} = \frac{\boldsymbol{r} - \boldsymbol{r}_g}{||\boldsymbol{r} - \boldsymbol{r}_g||} \ . \tag{48}$$

This tensor is symmetric and traceless. Since the cell surface is represented by triangular meshes, the formula must be adapted to work with triangles. We attribute a homogeneous signal to each triangle $t_i$ of the mesh, equal to the average of the signal of its three vertices. Eq. 48 then becomes:

$$Q_{\alpha\beta} = \frac{\sum_{i=1}^{N_t} q_{\alpha\beta}^{(i)} s(t_i)}{\sum_{i=1}^{N_t} A_i s(t_i)} \text{ with } q_{\alpha\beta}^{(i)} = \int_{t_i} dS \left(\overline{r}_\alpha \overline{r}_\beta - \frac{1}{3}\delta_{\alpha\beta}(\overline{r}_\gamma \overline{r}_\gamma)\right) \ , \tag{49}$$

where we have used the same conventions as in Eq. 37.

#### 5.8 Correlation of myosin nematic tensor with interface modes

We now wish to determine whether a correlation exists between the orientation of the myosin pattern described by $\boldsymbol{Q}$ and the direction of the saddle-node mode of the cell-cell interface, represented by $\boldsymbol{S}_{12}$ defined in Eq. 25. In the same spirit as in Eq. 43, we define a correlation operator $\overline{C}$ operating on two tensors $\boldsymbol{A}$, $\boldsymbol{B}$:

$$\overline{C}(\boldsymbol{A}, \boldsymbol{B}) = \frac{\sum_{\alpha,\beta} A_{\alpha\beta} B_{\alpha\beta}}{||\boldsymbol{A}|| \cdot ||\boldsymbol{B}||} \ , \tag{50}$$

where we use the norm of a tensor defined as:

$$||\boldsymbol{A}|| = \left( \sum_{\alpha,\beta} A_{\alpha\beta} \right)^{1/2} . \tag{51}$$

Given a vector $\boldsymbol{S}_{12}$ for the orientation of the saddle-node deformation mode, a nematic tensor can be defined according to:

$$Q^{S12}_{\alpha\beta} = S_{12\alpha} S_{12\beta} - \frac{1}{3} S_{12\gamma} S_{12\gamma} \delta_{\alpha\beta} . \tag{52}$$

One can then calculate a correlation between the cortical myosin nematic tensor and the orientation of the saddle-node deformation mode, according to:

$$\overline{C}(\boldsymbol{Q}^1 - \boldsymbol{Q}^2, \boldsymbol{Q}^{S12}) , \tag{53}$$

which is invariant by the exchange of cell indices $1 \leftrightarrow 2$. This correlation function is used in Fig. 4l.

### 6 Cylindrical/planar projections of basal/interfacial signal

In Ext. Fig. 3 we generate planar maps of the E-cadherin signal on the cell-cell interface as well as cylindrical maps of the myosin signal of the basal side of cells. For each timepoint, we created maps based on the coordinates of the vertices of the segmented cell meshes. As discussed above, the interface vertices as well as the basal vertices can be obtained from the segmented meshes. It is also possible to associate to each of these vertices an intensity signal.

#### 6.1 Alignment of maps using the rotation vector

We wish to project the fluorescent signals on the plane perpendicular to the $\boldsymbol{N}_{12}$ axis (interface map, E-cadherin) or on a cylinder around the $\boldsymbol{r}_{12}$ axis (basal map, myosin), as illustrated on Ext. Fig. 4f. Since the doublet rotates between time points, the reference frames must be rotated accordingly to prevent introducing a spurious rotation around the $\boldsymbol{r}_{12}$ axis. Therefore, for each time series, we compute the rotation matrices describing the rotation around the normalised rotation vector $\boldsymbol{\omega}_n = \frac{\boldsymbol{\omega}(t)}{||\boldsymbol{\omega}(t)||}$ between time $t$ and $t + \Delta t$:

$$\mathbf{R}(t) = \begin{bmatrix} \cos\beta + \omega_{nx}^2 (1 - \cos\beta) & \omega_{nx}\omega_{ny} (1 - \cos\beta) - \omega_{nz} \sin\beta & \omega_{nx}\omega_{nz} (1 - \cos\beta) + \omega_{ny} \sin\beta \\ \omega_{ny}\omega_{nx} (1 - \cos\beta) + \omega_{nz} \sin\beta & \cos\beta + \omega_{ny}^2 (1 - \cos\beta) & \omega_{ny}\omega_{nz} (1 - \cos\beta) - \omega_{nx} \sin\beta \\ \omega_{nz}\omega_{nx} (1 - \cos\beta) - \omega_{ny} \sin\beta & \omega_{nz}\omega_{ny} (1 - \cos\beta) + \omega_{nx} \sin\beta & \cos\beta + \omega_{nz}^2 (1 - \cos\beta) \end{bmatrix} , \tag{54}$$

where $\beta = \arccos(\boldsymbol{u}_{12}(t) \cdot \boldsymbol{u}_{12}(t + \Delta t))$ is the angle of rotation and $\boldsymbol{u}_{12}$ is the vector defined in Eq. 3.

#### 6.2 Basal cylindrical maps

In order to generate a cylindrical map of the combined basal vertices of the two cells, we construct a reference frame as:

$$\begin{aligned} \boldsymbol{e}_x^{\text{proj}} &= \boldsymbol{e}_{\text{rot}} - \left(\boldsymbol{e}_{\text{rot}} \cdot \frac{\boldsymbol{r}_{12}}{||\boldsymbol{r}_{12}||}\right) \frac{\boldsymbol{r}_{12}}{||\boldsymbol{r}_{12}||} \ , \\ (\boldsymbol{e}_x, \boldsymbol{e}_y, \boldsymbol{e}_z) &= \left(\frac{\boldsymbol{e}_x^{\text{proj}}}{||\boldsymbol{e}_x^{\text{proj}}||}, \boldsymbol{e}_z \times \boldsymbol{e}_x, \frac{\boldsymbol{r}_{12}}{||\boldsymbol{r}_{12}||}\right) \ , \end{aligned} \tag{55}$$

where $\boldsymbol{e}_{\text{rot}}(t+\Delta t) = \mathbf{R}(t)\boldsymbol{e}_{\text{rot}}(t)$ and $\boldsymbol{e}_{\text{rot}}(t=0)$ is a unit vector which can be adjusted to obtain different views of the cylindrical map.

We then use cylindrical coordinates to define a set of coordinates $(z, \phi)$ associated to each basal vertex $j$ of the cell $i$:

$$\boldsymbol{r}_j - \boldsymbol{r}_g^{(b)} = ||\boldsymbol{r}_j - \boldsymbol{r}_g^{(b)}|| \cos(\phi)\boldsymbol{e}_x + ||\boldsymbol{r}_j - \boldsymbol{r}_g^{(b)}|| \sin(\phi)\boldsymbol{e}_y + z\boldsymbol{e}_z \ , \tag{56}$$

with $\boldsymbol{r}_g^{(b)}$ the center of mass of the basal vertices of both cells in the doublet. We make equally spaced $z$ and $\phi$ bins and we average the myosin signal of the cells for each timepoint in the bins to finally construct a cylindrical map. Several examples can be found in Ext. Fig. 3d.

#### 6.3 Interfacial planar maps

In order to generate a planar map of the interface, we use the outcome of the interface shape analysis pipeline (section 4), which consists of a reference frame of the interface $(\boldsymbol{e}_X, \boldsymbol{e}_Y, \boldsymbol{N}_{12})$, a set of coordinates of interface vertices $\{(X_i, Y_i, H_i)$ for $i \in \{1, ..., N\}\}$ in this reference frame, and a polynomial $P(X,Y)$ fitting the interface. We use the fitted profile of the interface to generate a grid of points of coordinates $\{(X_i^{(\text{grid})}, Y_i^{(\text{grid})}, Z_i^{(\text{grid})} = P(X_i^{(\text{grid})}, Y_i^{(\text{grid})}))$ for $i \in \{1, ..., n_p^2\}\}$ defined as on Ext. Fig. 4g. The points are regularly spaced on a $(150 \times 150)$ grid, and are selected based on the 2D convex hull of the points $\{X_i, Y_i\}$. We transform these interface-based coordinates back to the 3D coordinates of the image:

$$\boldsymbol{r}_i^{(\text{grid})} = X_i^{(\text{grid})}\boldsymbol{e}_X + Y_i^{(\text{grid})}\boldsymbol{e}_Y + Z_i^{(\text{grid})}\boldsymbol{N}_{12} \ . \tag{57}$$

We extract the E-cadherin signal on these vertices. Finally, in order to generate the interfacial maps, we define rotated reference frames:

$$\begin{aligned} \boldsymbol{e}_x^{\text{proj}} =& \boldsymbol{e}_{\text{rot}} - \left(\boldsymbol{e}_{\text{rot}} \cdot \frac{\boldsymbol{N}_{12}}{||\boldsymbol{N}_{12}||}\right) \frac{\boldsymbol{N}_{12}}{||\boldsymbol{N}_{12}||} \ , \\ (\boldsymbol{e}_X^{\text{proj}}, \boldsymbol{e}_Y^{\text{proj}}, \boldsymbol{e}_Z^{\text{proj}}) =& \left(\frac{\boldsymbol{e}_x^{\text{proj}}}{||\boldsymbol{e}_x^{\text{proj}}||}, \boldsymbol{e}_Z^{\text{proj}} \times \boldsymbol{e}_X^{\text{proj}}, \frac{\boldsymbol{N}_{12}}{||\boldsymbol{N}_{12}||}\right) \ , \end{aligned} \tag{58}$$

where $\boldsymbol{e}_{\text{rot}}(t+\Delta t) = \mathbf{R}(t)\boldsymbol{e}_{\text{rot}}(t)$ and $\boldsymbol{e}_{\text{rot}}(t=0) = \boldsymbol{e}_X(t=0)$. We create new coordinates $(X^{\text{proj}}, Y^{\text{proj}})$ in which we display the interfacial maps:

$$\begin{aligned} X^{\text{proj}} &= \boldsymbol{r}^{(\text{grid})} \cdot \boldsymbol{e}_X^{\text{proj}} \ , \\ Y^{\text{proj}} &= \boldsymbol{r}^{(\text{grid})} \cdot \boldsymbol{e}_Y^{\text{proj}} \ . \end{aligned} \tag{59}$$

### 7 Interacting active surfaces simulations

#### 7.1 Description of the framework

In order to simulate a rotating doublet, we use the interacting active surfaces (IAS) framework developed by Torres-Sanchez et al [4] (Fig. 4d, Ext. Fig. 6f,g). The following section recapitulates the main features of IAS and additions that were brought to it in the present work. We use the same notation as in Ref. [4]. Cells are described as active surfaces, and the governing equations for the cell surface mechanics are discretised using a finite element method. In the limit of low Reynolds number, the mechanics of a single surface $S$ can be described by the following statement of the principle of virtual work:

$$\delta W = \int_S dS \left\{ \frac{1}{2}\hat{t}^{ij}\delta g_{ij} + \bar{m}^{ij}\delta C_{ij} - f^\alpha \delta X^\alpha \right\} = 0 \ , \tag{60}$$

where $\delta \boldsymbol{X}$ is an infinitesimal displacement of the surface, and $\delta g_{ij}$ and $\delta C_{ij}$ the associated infinitesimal variation of the metric and curvature tensors. This equation is equivalent to the statement of balance of linear and angular momentum at low Reynolds number. Constitutive equations are specified for the mechanical tensors $\hat{t}^{ij}$ and $\bar{m}^{ij}$. We denote by $\boldsymbol{v}$ the velocity field on the cell surface, which is assumed be an active viscous layer, with a bending rigidity:

$$\begin{aligned} \hat{t}^{ij} &= 2\eta\nu^{ij} + \gamma(\boldsymbol{r})g^{ij} + \kappa C_k^{\ k}\left(\frac{1}{2}C_k^{\ k}g^{ij} - 2C^{ij}\right) \ , \\ \bar{m}^{ij} &= \kappa C_k^{\ k} g^{ij} \ , \end{aligned} \tag{61}$$

where $\nu^{ij}$ is the strain rate tensor on the surface:

$$\nu^{ij} = \frac{1}{2}\left(\nabla^i v^j + \nabla^j v^i + 2C^{ij}v_n\right) \ , \tag{62}$$

with $\nabla^i$ the covariant derivative operator. Here, $\eta$ is the surface viscosity and $\gamma(\boldsymbol{r})$ is the surface tension of the cell, which we define in this study to be spatially modulated. The external force density acting on a single cell is given by:

$$f^\alpha = Pn^\alpha - \xi v^\alpha \ , \tag{63}$$

where $\xi$ is a friction coefficient with the outside medium and the pressure $P$ is adjusted to impose a value of the cell volume.

#### 7.2 Cell-cell adhesion

The previous equations are modified by the presence of an adhesion potential $\phi(|\boldsymbol{X}_I - \boldsymbol{X}_J|)$ operating between pair of points of cell $I$ and cell $J$. Each cell interaction with a cell $J$ is contributing an additional external force density on cell $I$, and an additional isotropic tension to cell $I$:

$$\begin{aligned} \boldsymbol{f}_{IJ} &= -\int_{S_J} dS_J \phi'(|\boldsymbol{X}_I - \boldsymbol{X}_J|)\frac{\boldsymbol{X}_I - \boldsymbol{X}_J}{|\boldsymbol{X}_I - \boldsymbol{X}_J|} \ , \\ t_{IJ}^{ij} &= \int_{S_J} dS_J \phi(|\boldsymbol{X}_I - \boldsymbol{X}_J|) g_I^{ij} \ . \end{aligned} \tag{64}$$

The potential of interaction is of the following form:

$$\phi(r) = D\left\{\left[1 - \exp\left(\frac{r_{min} - r}{l}\right)\right]^2 - 1\right\} w(r, r_{min}, r_{min} + 3l) \ , \tag{65}$$

with the cut-off function $w$ given by:

$$w(r, r_1, r_2) = \begin{cases} 1 & \text{if } r \leq r_1 \\ \left(\frac{r_2-r}{r_2-r_1}\right)^3 \left[6\left(\frac{r_2-r}{r_2-r_1}\right)^2 - 15\left(\frac{r_2-r}{r_2-r_1}\right) + 10\right] & \text{if } r_1 < r < r_2 \\ 0 & \text{if } r \geq r_2 \end{cases} \quad (66)$$

This potential has a minimum at $r = r_{min}$ with minimum value $-D$, and is also peaked around its minimum with characteristic length $l$. It goes to zero exactly at a distance $r_{min} + 3l$.

#### 7.3 Simulation setup and dimensionless parameters

We simulate a doublet of two cells 1 and 2 in contact. Each cell $i$ has a tension profile $\gamma_i(\boldsymbol{r})$:

$$\gamma_i(\boldsymbol{r}) = \gamma_i^0 + \gamma_i^m(\boldsymbol{r}) \ , \quad (67)$$

where $\gamma_i^0$ is the part of the surface tension (assumed to be spatially homogeneous) not linked to the myosin, and $\gamma_i^m(\boldsymbol{r})$ is the tension created by the myosin which is modulated around a unit polarity vector $\boldsymbol{P}_i$:

$$\gamma_i^m(\boldsymbol{r}) = \gamma_i^{m0} + \Delta\gamma_i \left( \frac{b_i \exp\left(b_i \boldsymbol{P}_i \cdot \frac{\boldsymbol{r}}{||\boldsymbol{r}||}\right) - \sinh(b_i)}{b_i \exp(b_i) - \sinh(b_i)} \right) \ . \quad (68)$$

Here, $\gamma_i^{m0}$ is a baseline tension and $\Delta\gamma_i$ is the amplitude of the tension modulation. The parameter $b_i$ represents how broad the tension modification around the point indicated by $\boldsymbol{P}_i$ is (Ext. Fig. 6a). For $b \to 0$ we reach the following tension modulation:

$$\gamma_i(\boldsymbol{r}) = \gamma_i^0 + \gamma_i^{m0} + \Delta\gamma_i \boldsymbol{P}_i \cdot \frac{\boldsymbol{r}}{||\boldsymbol{r}||} \ . \quad (69)$$

Then the size of the spot becomes smaller as $b$ increases (Ext. Fig. 6a,c). The maximum tension is $\gamma_i^0 + \gamma_i^{m0} + \Delta\gamma_i$ and the average tension on a spherical cell is $\gamma_i^0 + \gamma_i^{m0}$. This average tension is noted $\gamma_i^a$ in the following. We fix the angle $\theta_p$ between the cell polarity vector $\boldsymbol{P}_i$ of cell $i$ and the vector $\boldsymbol{r}_{ij}$ joining the center of mass of cell $i$ to the center of mass of cell $j$, using the following formulas:

$$\begin{aligned} \boldsymbol{P}_1 &= \cos(\theta_p)\frac{\boldsymbol{r}_{12}}{||\boldsymbol{r}_{12}||} + \sin(\theta_p)\boldsymbol{e}_z \times \frac{\boldsymbol{r}_{12}}{||\boldsymbol{r}_{12}||} \text{ with } \boldsymbol{e}_z = (0,0,1) \ , \\ \boldsymbol{P}_2 &= \cos(\theta_p)\frac{\boldsymbol{r}_{21}}{||\boldsymbol{r}_{21}||} + \sin(\theta_p)\boldsymbol{e}_z \times \frac{\boldsymbol{r}_{21}}{||\boldsymbol{r}_{21}||} \ . \end{aligned} \quad (70)$$

The doublet is initialized at $t = 0$ in such a way that $\boldsymbol{r}_{12} \cdot \boldsymbol{e}_z = 0$, which ensures that the rotation will occur around the axis defined by $\boldsymbol{e}_z$. In order to explore more complex tension profiles, we may add a second-order term $\gamma_{\text{nem},i}(\boldsymbol{r})$ to the tension profile $\gamma_i(\boldsymbol{r})$ defined in Eq. 67:

$$\gamma_{\text{nem},i}(\boldsymbol{r}) = \Delta\gamma_{n,i} \left( \left( \boldsymbol{P}_i \cdot \frac{\boldsymbol{r}}{||\boldsymbol{r}||} \right)^2 - \frac{1}{3} \right) \ . \quad (71)$$

On a spherical cell this term has a zero average. We make the system dimensionless by choosing units of time, space, and mass as follows:

$$T = \frac{\eta}{\gamma_a}, \ L = \left(\frac{3V}{4\pi}\right)^{1/3} = R, \ M = \frac{\eta^2}{\gamma_a} \text{ with } \gamma_a = \frac{1}{2}(\gamma_1^a + \gamma_2^a) \ , \quad (72)$$

with $V$ the volume of cell $i$, assumed identical to the volume of cell $j$. According to this choice, the remaining dimensionless parameters describe the phase space of the problem:

$$\frac{\gamma_1^a}{\gamma_2^a}, \frac{\Delta\gamma_1}{\gamma_1^a}, \frac{\Delta\gamma_2}{\gamma_2^a}, \frac{\Delta\gamma_{n,1}}{\gamma_1^a}, \frac{\Delta\gamma_{n,2}}{\gamma_2^a}, b_1, b_2, \theta_p, \frac{\kappa}{\gamma_a R^2}, \frac{\xi R^2}{\eta}, \frac{DR^2}{\gamma_a}, \frac{r_{min}}{R}, \frac{l}{R} \; . \tag{73}$$

#### 7.4 Comparison of simulations with experiments

Here we discuss the relationship between active tension profiles used in simulation and experimentally obtained cortical myosin profiles. We assume that the fluorescence intensity $I(\boldsymbol{r})$ of myosin is proportional to the myosin generated tension $\gamma^m(\boldsymbol{r})$:

$$\gamma^m(\boldsymbol{r}) = \nu I(\boldsymbol{r}) \; . \tag{74}$$

We quantify the contrast in the experimental myosin signal using the standard deviation divided by the average:

$$\frac{\sigma_I}{\langle I \rangle} = \frac{\sqrt{\langle I^2 \rangle - \langle I \rangle^2}}{\langle I \rangle} = \sqrt{\frac{\langle I^2 \rangle}{\langle I \rangle^2} - 1} \; , \tag{75}$$

where the averages $\langle ... \rangle$ are computed after projection on a unit sphere, like in section 5.3. In simulations, we compute the tension contrast similarly as $\sigma_\gamma / \langle \gamma \rangle$. Using Eq. 67 and Eq. 74, we get the following relation between the total tension $\gamma$ and the intensity:

$$\gamma = \gamma^0 + \nu I \; , \tag{76}$$

which leads to:

$$\frac{\sigma_I}{\langle I \rangle} = \frac{\sigma_{\gamma^m}}{\langle \gamma^m \rangle} = \left(1 + \frac{\gamma^0}{\nu \langle I \rangle}\right) \frac{\sigma_\gamma}{\langle \gamma \rangle} \tag{77}$$

This shows that in order to compare a simulation result to an experimental result, one should use $\sigma_{\gamma^m}/\langle \gamma^m \rangle$. However, because of the underlying and unmeasured tension $\gamma^0$, this requires to determine the value of the unknown scaling coefficient $\lambda = (1 + \gamma^0/(\nu \langle I \rangle))$. We determine the value of $\lambda$ by superimposing the experimental and simulated curves of $\sqrt{\langle H_{y-y}^2 \rangle}/R$, since the deflection of the interface is a purely geometric quantity that should be identical in an experiment and its equivalent simulation. This leads to $\lambda = 6$, which would mean that the average unmeasured tension $\gamma^0$ is typically 5 times higher than the average measured tension $\nu \langle I \rangle$. We use this value of $\lambda$ on Fig. 4h and Fig. 4i to compute $\sigma_{\gamma^m}/\langle \gamma^m \rangle$.

We now consider two cells 1 and 2 of a doublet with two homogeneous, but possibly distinct surface tensions $\gamma_1 = \gamma_1^a = \gamma^0 + \gamma_1^m$ and $\gamma_2 = \gamma_2^a = \gamma^0 + \gamma_2^m$. We would like to compare this situation with the shape of a doublet with average myosin fluorescence intensity $\langle I_1 \rangle$ and $\langle I_2 \rangle$. Here we assume that the unmeasured tension $\gamma^0$ is the same for both cells, as well as the conversion coefficient $\nu$. Eq. 74 then shows that we have:

$$\frac{\gamma_1^m - \gamma_2^m}{\gamma_1^m + \gamma_2^m} = \frac{\langle I_1 \rangle - \langle I_2 \rangle}{\langle I_1 \rangle + \langle I_2 \rangle} \; . \tag{78}$$

where the averaging operator $\langle \rangle$ has been defined in Eq. 47. Since $\lambda$ is defined in average on a population of cells, we can assume that:

$$\lambda = 1 + \frac{\gamma^0}{\nu \frac{\langle I_1 \rangle + \langle I_2 \rangle}{2}} = 1 + \frac{2\gamma^0}{\gamma_1^m + \gamma_2^m} \; . \tag{79}$$

From this we obtain that $\gamma_1^a + \gamma_2^a = 2\gamma_0 + \gamma_1^m + \gamma_2^m = \lambda(\gamma_1^m + \gamma_2^m)$, which means that the left hand side of Eq. 78 can be computed as:

$$\frac{\gamma_1^m - \gamma_2^m}{\gamma_1^m + \gamma_2^m} = \lambda \frac{\gamma_1^a - \gamma_2^a}{\gamma_1^a + \gamma_2^a} \ . \tag{80}$$

We use this formula to compare the experimental and simulated curves of the bowl mode amplitudes in Fig. 4k.

#### 7.5 Parameter values used in simulations

The parameters $r_{min}/R$ and $l/R$ are chosen so that for a cell of typical radius 5$\mu$m we have $r_{min}$ = 600nm and $l$ = 200nm. The value $r_{min}$ sets the distance between the cortices of the interacting cells. The width of an adherens junction is roughly 20nm [5],and a typical cortex thickness is 200nm [7], bringing the total distance between the middle of the two cortices to $\sim$ 220nm. Here we use a larger value, but with the right order of magnitude. We note that it is important for numerical stability that the parameters $r_{\min}$, $l$ are not small compared to the typical length scale of the triangle mesh used for the simulation. Here we use meshes with 20480 triangles with a typical length (square root of area) of $0.03/R$, and we have chosen a value of $l/R = 0.04$ and $r_{min}/R = 0.12$.

For all the simulations, we set the angle $\theta_p$ of the polarity to $\pi/2$, which is close to what is experimentally measured (Fig. 3g). We also fixed the adhesion strength to $DR^2/\gamma_a = 38.9$. The value of the adhesion strength (relative to the surface tension) sets the equilibrium area of contact between the two doublets and is chosen here so that the doublet has an overall spherical shape when it is not rotating (see Fig. 4j, $D_{\infty h}$).

For the bending rigidity we operate around the value of $\kappa/(\gamma_a R^2) = 10^{-2}$ which, for a typical cortical tension $\gamma_a \approx 10^{-4} N.m^{-1}$ [8], corresponds to $\kappa \approx 6 \times 10^3 k_B T$. This is compatible with a typical cortical Young modulus $E \approx 1$ kPa, and a cortex thickness $h \approx 200$ nm, which lead to an estimated bending rigidity $Eh^3 \approx 2 \times 10^3 k_B T$.

For $\xi R^2/\eta$ we choose a default value of 10, which corresponds to a typical hydrodynamic length $\sqrt{\eta/\xi} \approx 1\mu m$. This quantity can be expected to vary a lot depending on the friction with the outside medium. It has been estimated in *C.elegans* zygotes to be around 14$\mu$m [9]. The values of the remaining parameters are listed in Table 3.

Simulations can be run with the normalized time $t\gamma_a/\eta$ and space coordinates normalized by $R$. In Fig. 4h, we obtain rotation velocities comparable to experiments by setting $\eta/\gamma_a$ = 1min, and in Ext. Fig. 6f we then obtain cortical flow velocities in $\mu$m/min by setting in addition $R = 5\mu$m.

All simulations are started from a doublet initially at rest at $t\gamma_a/\eta = 0$ (steady-state configuration of Fig. 4j) and are run until the rotational velocity reaches a steady-state. We typically see small oscillations in the interface deflection (Ext. Fig. 6e). For the laser ablation simulation (Fig. 5d,e), the tension modulations of both cells are turned to 0 at a time $t\gamma_a/\eta = 175$ corresponding to the steady-state of the reference simulation (Fig. 4e).

For the optogenetic simulation (Fig. 5f,g,h), we also start from the steady-state of the reference simulation and we introduce an additional tension modulation on cell 1:

$$\gamma_{\text{opto}}(\boldsymbol{r}) = \Delta\gamma_{\text{opto}} \left( \frac{b_{\text{opto}} \exp\left( b_{\text{opto}} \boldsymbol{P}_{\text{opto}} \cdot \frac{\boldsymbol{r}}{||\boldsymbol{r}||} \right) - \sinh(b_{\text{opto}})}{b_{\text{opto}} \exp(b_{\text{opto}}) - \sinh(b_{\text{opto}})} \right) \ , \tag{81}$$

| Figure | $\frac{\gamma_1^0}{\gamma_2^0}$ | $\frac{\Delta\gamma_1}{\gamma_1^0}$ | $\frac{\Delta\gamma_2}{\gamma_2^0}$ | $\frac{\Delta\gamma_{n,1}}{\gamma_1^0}$ | $\frac{\Delta\gamma_{n,2}}{\gamma_2^0}$ | $b_1$ | $b_2$ | $\frac{\kappa}{\gamma_a R^2}$ | $\frac{\xi R^2}{\eta}$ |
|---|---|---|---|---|---|---|---|---|---|
| 4e,f,g, Ext.Fig.6e,f | 1 | 0.3 | 0.3 | 0 | 0 | 0 | 0 | $10^{-2}$ | 10 |
| 4h,i | 1 | x | x | 0 | 0 | 0 | 0 | $10^{-2}$ | 10 |
| 4j:$D_{\infty h}$ | 1 | 0 | 0 | 0 | 0 | 0 | 0 | $10^{-2}$ | 10 |
| 4j:$C_{2h}$ | 1 | 0.3 | 0.3 | 0 | 0 | 0 | 0 | $10^{-2}$ | 10 |
| 4j:$C_{\infty v}$ | 1.171 | 0 | 0 | 0 | 0 | 0 | 0 | $10^{-2}$ | 10 |
| 4j:$C_{2v}$ | 1 | 0 | 0 | 0.1 | 0 | 0 | 0 | $10^{-2}$ | 10 |
| 4j:$C_s$ | 1.041 | 0.288 | 0.3 | 0.072 | 0 | 3 | 3 | $9.8 \cdot 10^{-3}$ | 10 |
| 4k | x | 0 | 0 | 0 | 0 | 0 | 0 | $10^{-2}$ | 10 |
| 5d,e | 1 | 0.3 | 0.3 | 0 | 0 | 0 | 0 | $10^{-2}$ | 10 |
| 5f,g,h | 1 | 0.3 | 0.3 | 0 | 0 | 0 | 0 | $10^{-2}$ | 10 |
| Ext.Fig.6d | 1 | 0.3 | 0.3 | 0 | 0 | x | x | $10^{-2}$ | 10 |
| Ext.Fig.6h | 1 | 0.3 | 0.3 | 0 | 0 | 0 | 0 | x | x |

**Supplementary Table 3:** Parameter values used in simulations. An “x” is shown in the table above when the parameter values are directly indicated on the corresponding figures.

with $\mathbf{P}_{\text{opto}}$ a unit vector. We choose $\Delta\gamma_{\text{opto}}/\gamma_a = 0.3$ and $b_{\text{opto}} = 4$. The position of the spot is adjusted to follow the rotation of doublet (which is around the axis $\boldsymbol{e}_z = (0, 0, 1)$) using the following formula:

$$\begin{aligned} \boldsymbol{P}_1 &= \cos\left(\frac{\pi}{2}\right)\frac{\boldsymbol{r}_{12}}{||\boldsymbol{r}_{12}||} + \sin\left(\frac{\pi}{2}\right)\boldsymbol{e}_y \\ \boldsymbol{P}_{\text{opto}} &= \cos\left(-\frac{\pi}{1.8}\right)\frac{\boldsymbol{r}_{12}}{||\boldsymbol{r}_{12}||} + \sin\left(-\frac{\pi}{1.8}\right)\boldsymbol{e}_y \\ \text{with } \boldsymbol{e}_y &= \frac{\boldsymbol{e}_z \times \boldsymbol{r}_{12}}{||\boldsymbol{e}_z \times \boldsymbol{r}_{12}||} . \end{aligned} \tag{82}$$

#### 7.6 Effect of tension spot size on interface deformation

As is seen on Fig. 4j (second row), and on supplementary table 1, the yin-yang and three-fold modes are similar in the sense that they both possess the necessary symmetries to be observed in a typical rotating doublet simulation like the one of Fig. 4e. However, for a broad tension profile ($b = 0$), only the yin-yang mode is observed (Fig. 4f). Interestingly, on Ext. Fig. 6d we show that by reducing the size of the active tension spot, one can generate rotating doublets with a three-fold deformation mode, in addition to the yin-yang mode. For $b > 3$, the three-fold mode even dominates in amplitude. We chose a range for the $b$ parameter based on the experimental values found when fitting a profile of the type of Eq.68 to the experimental myosin signal at different individual time points:

$$I = \langle I \rangle + \Delta I \left( \frac{b \exp\left(b\boldsymbol{P} \cdot \frac{\boldsymbol{r}}{||\boldsymbol{r}||}\right) - \sinh(b)}{b\exp(b) - \sinh(b)} \right) , \tag{83}$$

with $\boldsymbol{P}$ being the normalised polarity vector defined in Eq.37. The fitted values of $b$ and $\Delta I/\langle I \rangle$ are shown on Ext. Fig. 6b.

#### 7.7 Effect of bending rigidity and friction

On Ext. Fig. 6h, we test the effect of the dimensionless bending rigidity $\kappa/(\gamma_a R^2)$ and the dimensionless friction coefficient $\xi R^2/\eta$ on the rotational velocity and the interface deflection.We see that the bending rigidity acts on both observables, but mostly acts on the interface deflection. For increasing rigidity, the interface is less deflected. The friction coefficient, on the opposite, acts mostly on the rotational velocity. A higher friction coefficient leads to a slower rotation.
